# Declining juvenile survival of Adélie penguins in Antarctica

**DOI:** 10.1101/2024.08.23.609289

**Authors:** Téo Barracho, Gaël Bardon, Rémi Choquet, Aymeric Houstin, Alexander Winterl, Michaël Beaulieu, Pierrick Blanchard, Cindy C. Cornet, Robin Cristofari, Thierry Raclot, Sebastian Richter, Nicolas Chatelain, Julien Courtecuisse, Matthieu Brucker, Daniel P. Zitterbart, Nicolas Lecomte, Céline Le Bohec

## Abstract

As summer sea ice around Antarctica reaches modern lows, quantifying the demographic response of polar species to such environmental changes becomes critical. To achieve this, synthesizing results across species’ ranges and elucidating the environmental factors driving population dynamics are key. Adélie penguins are considered reliable indicators of changes in Antarctica but the processes through which sea ice and other environmental factors shape their population dynamics are still unclear, especially for critical age groups such as juveniles. Using a 17-year dataset of Adélie penguins electronically tagged in Adélie Land (Antarctica), we found that juvenile survival probability was most impacted by sea ice concentration near their natal colony right after fledging, with lower ice concentrations detrimental to survival. Importantly, we found that juvenile survival declined by 32% from 2007 to 2020, mirroring trends at other distant colonies. The emergence of similar patterns at opposite ends of the continent may be an early signal for shifts in population trends expected from climate change.

## Introduction

Antarctica is at the center of large and rapid environmental shifts, with both accelerating climate changes^1–3^ and increasing exploitation pressure by fisheries^4^. Quantifying the response of animal populations to such environmental forcing is essential to determine their potential dynamics and persistence^5,6^. Yet, our understanding of how Antarctic species respond to such changes is still limited. Typically, the spatially and temporally heterogeneous dynamics of climate^7,8^ and of environmental parameters such as sea ice^9^ across species distributions makes identifying drivers of any demographic rate challenging^10–12^. This hampers our ability to anticipate species response to environmental changes, especially over scales relevant to ecosystem management^13,14^.

The Adélie penguin (*Pygoscelis adeliae*), an iconic Antarctic species, exemplifies this challenge particularly well. Early observations of contrasting population trends between the Ross Sea and the Antarctic Peninsula in response to sea ice declines have fostered a population dynamics model, where abundance is maximal at intermediate sea ice concentrations^15,16^. This model has been instrumental in projecting the response of Adélie penguins to climate variability across large spatial and temporal scales^17–19^. However, more localized and shorter-term predictions are still hindered by the inherent variability and noise in abundance time series, which most population studies are constrained to rely on^19–21^. Precise quantification of key demographic rates (e.g. juvenile survival) is critical for elucidating population dynamics driven by complex environmental interactions^22,23^. The need for such an approach is twofold; it can provide short-term feedback for ecological management at the local scale and be integrated at larger spatial scales as more studies become available across the species range^24–27^.

In long-lived species, adult survival typically holds the greatest potential to influence population dynamics, but actual population growth rates can often be driven by other demographic rates such as fecundity or juvenile survival^28^. In long-lived Adélie penguins, juvenile survival indeed appears to play a prominent role in driving population fluctuations. For example, Adélie penguin’s abundance (as measured by the number of breeding pairs) correlates with sea ice conditions with a 5-years lag^29^; which is consistent with effects of sea ice on juvenile survival during their first months at sea^30^. In the Antarctic Peninsula^31,32^ and East Antarctica^33^, population size decreased when juvenile survival also did, further emphasizing its central role in shaping overall population trends. Despite such findings, we still lack clear evidence linking juvenile survival to environmental parameters known to affect population dynamics, such as sea ice concentration^13^. Although adult survival has been convincingly linked to environmental variables (e.g., Southern Oscillation Index, winter sea ice concentration) in several populations around Antarctica^31,34–37^, the drivers of juvenile survival remain unclear^31,33,36^. Given the importance of this demographic parameter, evaluating its trends across the species’ range and elucidating its environmental drivers are required to adequately predict the species’ response to future environmental changes^38^.

Here, we quantify juvenile survival probabilities for the first time in the Western Pacific Ocean sector (90°E to 160°E, Fig. S1) using a 17-year-long dataset of known-age, electronically tagged Adélie penguins from Pointe Géologie archipelago, Adélie Land. By investigating trends in survival for this population located thousands of kilometers away from other studied colonies and synthesizing all previously published studies on this vital rate, we provide a comparative picture of trends in Adélie penguin juvenile survival across Antarctica. To uncover the mechanisms behind variations in juvenile survival, we also investigate the drivers of juvenile survival at Pointe Géologie by linking survival probabilities to intrinsic and environmental parameters such as body mass at fledging and sea ice concentrations during their first winter at sea.

## Results

We used a multi-state capture-recapture framework accounting for short-range dispersion within a 10-km radius to model the survival probability of juvenile Adélie penguins (0-2 years old). Based on 17 years of tagging and detection data of 2807 individuals (2007-2023, see methods and Table S4), this model enabled us to quantify juvenile survival probabilities over 14 years (2007-2020). Over this period, juvenile survival estimates averaged 0.42 ± 0.18 and showed high inter-annual variability (CV = 44%, Fig. 2A). Notably, juvenile survival probabilities exhibited a negative temporal trend (Fig. 2A; Table 1, model 36, analysis of deviance: *p*-ANODEV = 0.016). The annual rate of change was of -2.5% (± 95%CI: -3.1, -2.0), translating into a 32% loss between 2007 and 2020. Juvenile survival has previously been quantified at 5 other locations around Antarctica^31,33,35,39,40^ (summarized in Fig. 2). Temporal variation was investigated at only two of them, and juvenile survival probabilities declined at both (Admiralty Bay, King George Island, Western Antarctic Peninsula, -1.3% per year for 1982-2000 ^31^; Béchervaise Island, Mac. Robertson Land, East Antarctica, -1.8% per year for 1992-2015 ^33^; Fig. 2).

**Fig. 1.**
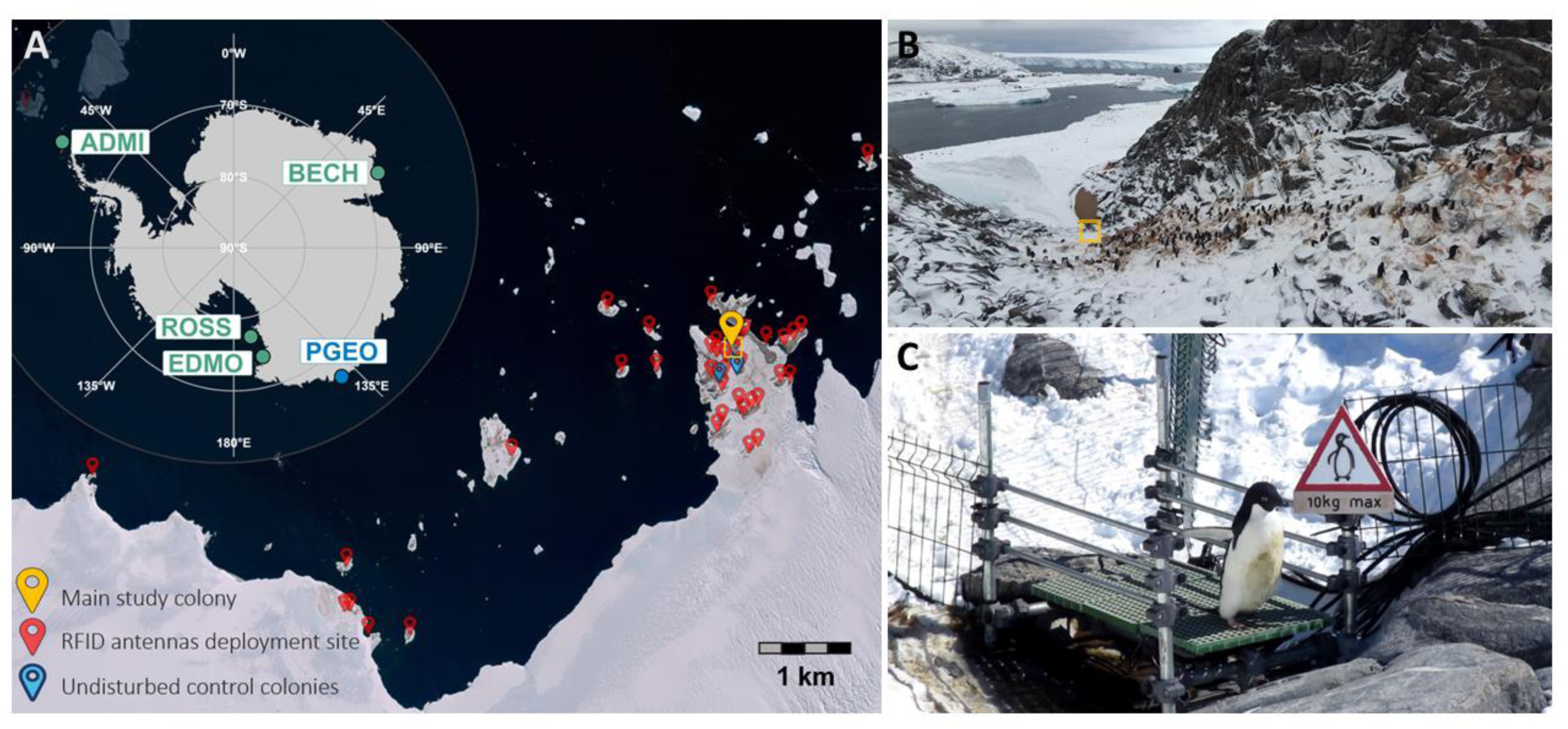
Overview of the Adélie penguin individual monitoring system operating in Pointe Géologie archipelago (Adélie Land) and location of similar monitoring sites across Antarctica still active as of 2024. **(A)** Main panel: Satellite imagery of Pointe Géologie archipelago (Pléiades Neo, Airbus DS 2021, 12/10/2021) with locations of the mobile RFID-antennas deployment sites (red), undisturbed control colonies (blue) and the main study colony (yellow). Top-left inset: location of study sites across Antarctica where longitudinal monitoring of marked Adélie penguins is still carried out (ADMI: Admiralty Bay, BECH: Béchervaise Island, EDMO: Edmonson Point, PGEO: Pointe Géologie, ROSS: Ross Island (4 monitored colonies including one on neighboring Beaufort Island)). **B)** Study colony in Pointe Géologie where all fledged chicks have been implanted with RFID-tags since 2007. **(C)** Penguins can only access or exit through two RFID-equipped passageways (only one shown here).

**Fig. 2.**
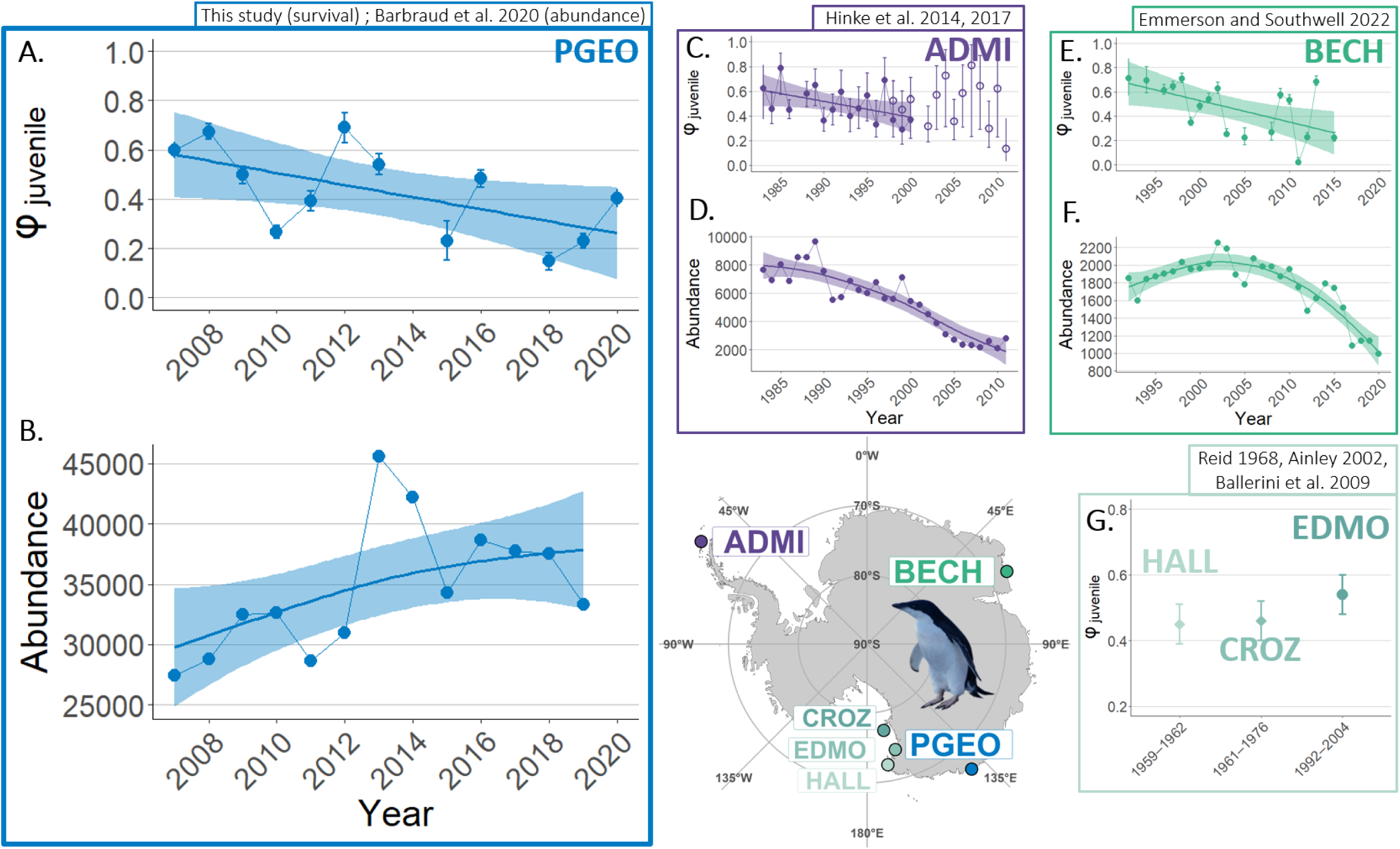
Summary of the six locations around Antarctica where Adélie penguin juvenile survival (ɸ, 0-2 years old) has been quantified. Among them, only three have investigated temporal variations. **A)** Survival of juvenile Adélie penguins from Pointe Géologie archipelago, Adélie Land (PGEO, electronic tagging, 2007-2020). The dots and intervals represent annual survival probabilities ± SE. The regression line and associated SE are from a linear regression (p = 0.036, R² = 0.37); **B)** Adélie penguin abundance (number of breeding pairs) at Pointe Géologie Archipelago (2007-2019, data reproduced from^74^); **C-D)** Juvenile survival (1982-2000 aluminum bands, 1998-2011 stainless steel bands) and abundance (1982-2011) at Admiralty Bay, King George Island, Western Antarctic Peninsula (ADMI, data reproduced from^31,32^); **E-F)** Juvenile survival (electronic tagging, 1992-2015) and abundance (1992-2020) at Béchervaise Island, Mac. Robertson Land (BECH, data reproduced from^33^); **G)** Only mean estimates were available for three populations in the Ross Sea: Cape Hallett, Victoria Land (HALL, metal bands, 1959-1962, data extracted from^39^), Cape Crozier, Ross Island (CROZ, aluminum bands, 1961-1976, data extracted from^40^), Edmondson Point, Victoria Land (EDMO, electronic tagging, 1994-2002, data extracted from^35^). Survival estimates from HALL and to a lesser extent CROZ should be compared to other estimates with caution because they were not computed using modern capture-recapture techniques accounting for imperfect detection as in the other studies. Juvenile survival time series were fitted with linear regressions when authors reported a linear trend (**A,C,E**). Abundance time series were fitted with Generalized Additive Models (GAM).

**Table 1.**
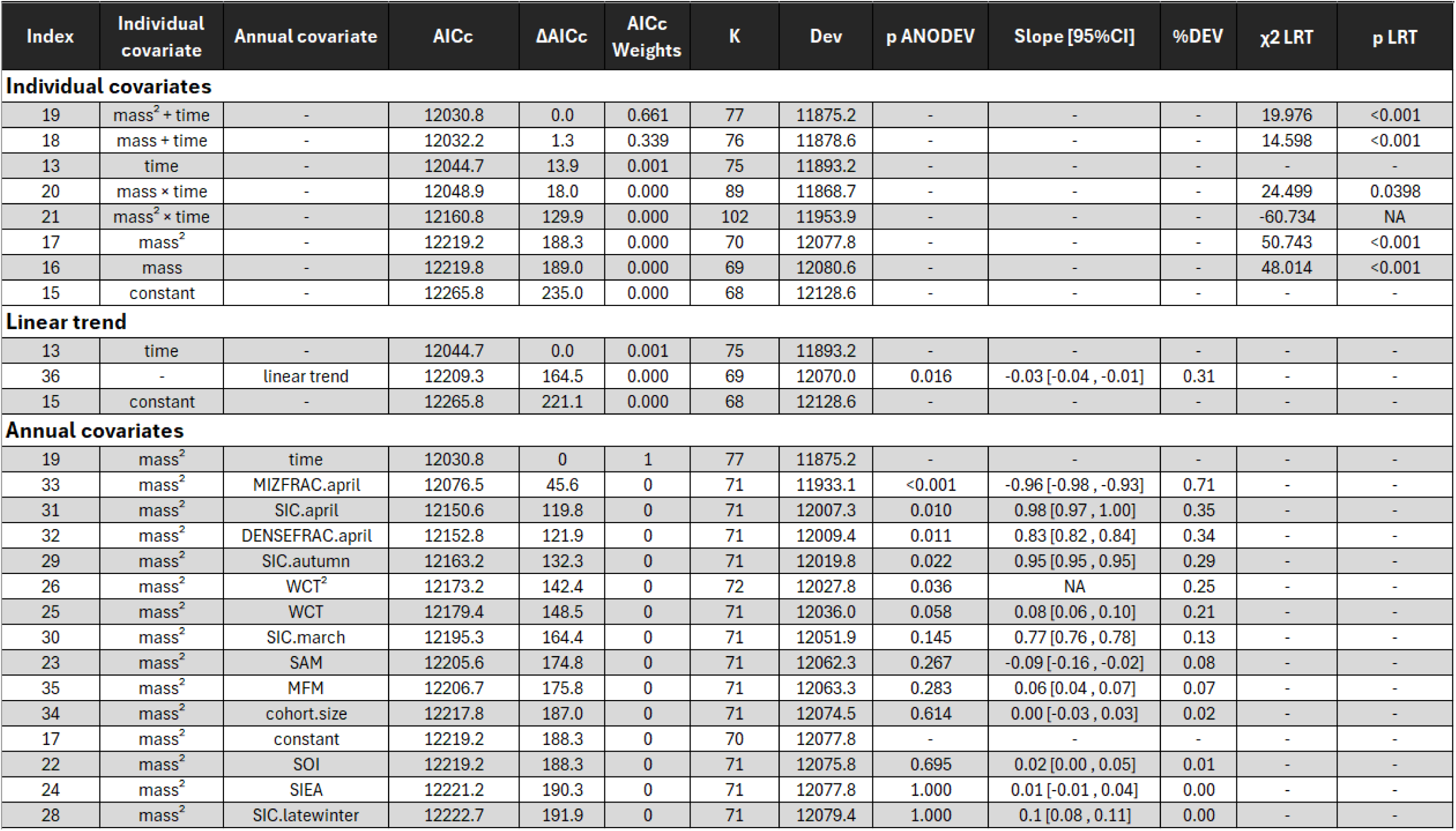
Comparison of the effect of covariates on Adélie penguin juvenile survival probability. Abbreviations: AICc, Akaike Information Criterion corrected for small sample sizes; ΔAICc, AICc difference between current model and time-dependant model; K, number of parameters; Dev, Model deviance; p ANODEV, p-value of the analysis of deviance; %DEV, percentage of the time-dependent model’s deviance explained by the inclusion of covariate^25^; LRT, Likelihood Ratio Test. Covariates abbreviations: see footnote and Table S2. Footnote Table 1: MIZFRAC.april (April fraction of total ice area between 135°E-145°E covered by the marginal ice zone (15-80% SIC)); DENSEFRAC.april (April fraction of total ice area between 135°E-145°E covered by dense ice (80-100% SIC)); SIC.april (April average SIC in area between 135°E-145°E); SIC.autumn (Mar-Apr average SIC in area between 135°E-145°E); WCT (Windchill temperatures for Dumont D’Urville in Apr-Sept); SAM (Southern Annular Mode, Mar-Sept); SIC.march (March average SIC in area between 135°E-145°E); SIC.latewinter (Aug-Sept average SIC in area between 110°E-135°E); SOI (Southern Oscillation Index, Mar-Sept); SIEA (Sea Ice Extent Anomalies, Mar-Sept), SIC.winter (May-Jul average SIC in area between 110°E-135°E).

| Index | Individual covariate | Annual covariate | AICc | $\Delta AICc$ | AICc Weights | K | Dev | p ANODEV | Slope [95%CI] | %DEV | $\chi^2$ LRT | p LRT |
| --- | --- | --- | --- | --- | --- | --- | --- | --- | --- | --- | --- | --- |
| **Individual covariates** |  |  |  |  |  |  |  |  |  |  |  |  |
| 19 | mass<sup>2</sup> + time | - | 12030.8 | 0.0 | 0.661 | 77 | 11875.2 | - | - | - | 19.976 | <0.001 |
| 18 | mass + time | - | 12032.2 | 1.3 | 0.339 | 76 | 11878.6 | - | - | - | 14.598 | <0.001 |
| 13 | time | - | 12044.7 | 13.9 | 0.001 | 75 | 11893.2 | - | - | - | - | - |
| 20 | mass × time | - | 12048.9 | 18.0 | 0.000 | 89 | 11868.7 | - | - | - | 24.499 | 0.0398 |
| 21 | mass<sup>2</sup> × time | - | 12160.8 | 129.9 | 0.000 | 102 | 11953.9 | - | - | - | -60.734 | NA |
| 17 | mass<sup>2</sup> | - | 12219.2 | 188.3 | 0.000 | 70 | 12077.8 | - | - | - | 50.743 | <0.001 |
| 16 | mass | - | 12219.8 | 189.0 | 0.000 | 69 | 12080.6 | - | - | - | 48.014 | <0.001 |
| 15 | constant | - | 12265.8 | 235.0 | 0.000 | 68 | 12128.6 | - | - | - | - | - |
| **Linear trend** |  |  |  |  |  |  |  |  |  |  |  |  |
| 13 | time | - | 12044.7 | 0.0 | 0.001 | 75 | 11893.2 | - | - | - | - | - |
| 36 | - | linear trend | 12209.3 | 164.5 | 0.000 | 69 | 12070.0 | 0.016 | -0.03 [-0.04, -0.01] | 0.31 | - | - |
| 15 | constant | - | 12265.8 | 221.1 | 0.000 | 68 | 12128.6 | - | - | - | - | - |
| **Annual covariates** |  |  |  |  |  |  |  |  |  |  |  |  |
| 19 | mass<sup>2</sup> | time | 12030.8 | 0 | 1 | 77 | 11875.2 | - | - | - | - | - |
| 33 | mass<sup>2</sup> | MIZFRAC.april | 12076.5 | 45.6 | 0 | 71 | 11933.1 | <0.001 | -0.96 [-0.98, -0.93] | 0.71 | - | - |
| 31 | mass<sup>2</sup> | SIC.april | 12150.6 | 119.8 | 0 | 71 | 12007.3 | 0.010 | 0.98 [0.97, 1.00] | 0.35 | - | - |
| 32 | mass<sup>2</sup> | DENSEFRAC.april | 12152.8 | 121.9 | 0 | 71 | 12009.4 | 0.011 | 0.83 [0.82, 0.84] | 0.34 | - | - |
| 29 | mass<sup>2</sup> | SIC.autumn | 12163.2 | 132.3 | 0 | 71 | 12019.8 | 0.022 | 0.95 [0.95, 0.95] | 0.29 | - | - |
| 26 | mass<sup>2</sup> | WCT<sup>2</sup> | 12173.2 | 142.4 | 0 | 72 | 12027.8 | 0.036 | NA | 0.25 | - | - |
| 25 | mass<sup>2</sup> | WCT | 12179.4 | 148.5 | 0 | 71 | 12036.0 | 0.058 | 0.08 [0.06, 0.10] | 0.21 | - | - |
| 30 | mass<sup>2</sup> | SIC.march | 12195.3 | 164.4 | 0 | 71 | 12051.9 | 0.145 | 0.77 [0.76, 0.78] | 0.13 | - | - |
| 23 | mass<sup>2</sup> | SAM | 12205.6 | 174.8 | 0 | 71 | 12062.3 | 0.267 | -0.09 [-0.16, -0.02] | 0.08 | - | - |
| 35 | mass<sup>2</sup> | MFM | 12206.7 | 175.8 | 0 | 71 | 12063.3 | 0.283 | 0.06 [0.04, 0.07] | 0.07 | - | - |
| 34 | mass<sup>2</sup> | cohort.size | 12217.8 | 187.0 | 0 | 71 | 12074.5 | 0.614 | 0.00 [-0.03, 0.03] | 0.02 | - | - |
| 17 | mass<sup>2</sup> | constant | 12219.2 | 188.3 | 0 | 70 | 12077.8 | - | - | - | - | - |
| 22 | mass<sup>2</sup> | SOI | 12219.2 | 188.3 | 0 | 71 | 12075.8 | 0.695 | 0.02 [0.00, 0.05] | 0.01 | - | - |
| 24 | mass<sup>2</sup> | SIEA | 12221.2 | 190.3 | 0 | 71 | 12077.8 | 1.000 | 0.01 [-0.01, 0.04] | 0.00 | - | - |
| 28 | mass<sup>2</sup> | SIC.latewinter | 12222.7 | 191.9 | 0 | 71 | 12079.4 | 1.000 | 0.1 [0.08, 0.11] | 0.00 | - | - |

To understand the drivers behind the decline observed at our study site, we used our previously developed model to test the effect of various intrinsic (mass at fledging, cohort size) and environmental covariates such as sea ice concentrations (SIC), windchill temperatures, and global climatic indices (Southern Oscillation Index, Southern Annular Mode, Table S2) on juvenile survival probabilities. Local and regional variables (Table S2) explained more of the variation in juvenile survival rates than global indices of climate variability (Table 1). Autumn sea ice concentrations (SIC.autumn) within 200 km of the natal colony (where juveniles are expected to be following fledging) correlated positively with juvenile survival rates (Fig. 3A, Table 1, model 29, %DEV = 28.6, slope ± 95%CI = 0.95 (0.94, 0.95)). We also found a quadratic effect of winter windchill temperatures (Table 1, model 26, %DEV = 24.7), with warmer temperatures associated with higher juvenile survival.

**Fig. 3.**
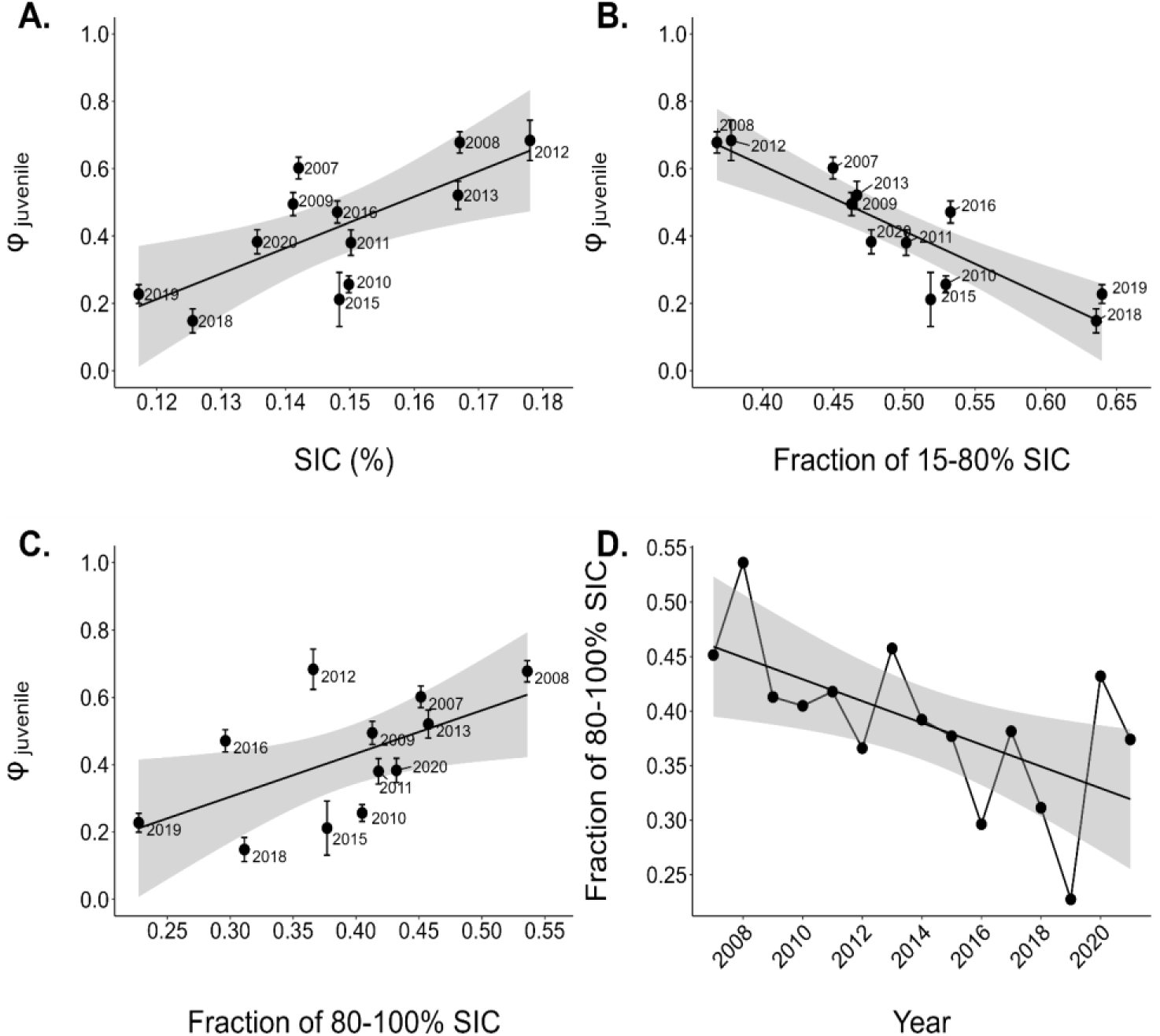
Relationships between the survival (ɸ) of juvenile Adélie penguins from Pointe Géologie archipelago (Adélie Land, East Antarctica) and several sea ice metrics within 200 kilometers of the natal colony for the month of April. **A)** Average sea ice concentration (SIC), **B)** Fraction of the total ice area (15-100% SIC) covered by the marginal ice zone (15-80% SIC), **C)** Fraction of total ice area (15-100% SIC) covered by dense ice (80-100% SIC). **D)** Time series (2007-2021) of the fraction of total ice area (15-100% SIC) covered by dense ice (80-100% SIC). Black thick lines and associated SE (gray shadow) on each panel are from linear regressions (respectively: p = 0.009, R² = 0.51; p < 0.001, R² = 0.79; p = 0.049, R² = 0.33; p = 0.016, R² = 0.37).

To better understand how SIC could affect juvenile survival at the onset of winter, we decomposed SIC.autumn into monthly data (SIC.march and SIC.april), and only SIC.april explained more than 20% DEV (Table 1, model 31, %DEV = 34.8). By decomposing SIC.april into the fraction of total ice area (15-100% SIC) covered by the marginal ice zone (15-80% SIC, MIZFRAC.april) and the fraction of total ice area covered by dense sea ice (80-100% SIC, DENSEFRAC.april), we then found that the amount of loose sea ice in the vicinity of the colony (MIZFRAC.april) was negatively related to juvenile survival rates (slope ± 95%CI = - 0.96 (- 0.98, - 0.93), Fig. 3B) and accounted for 71.4% of the temporal variation (Table 1, model 33). Conversely, the amount of dense sea ice (DENSEFRAC.april) was positively related to juvenile survival rates (slope ± 95%CI = 0.83 (0.82, 0.84), Fig. 3C) and accounted for 33.8% of the temporal variation (Table 1, model 32). This amount of April dense sea ice near the colony declined by 13% between 2007 and 2020 (linear model: F = 7.65, *p* = 0.016, R² = 0.37, Fig. 3D).

Chick body mass at tagging (fledging mass), an important driver of post-fledging survival in birds, averaged 3.57 ± 0.62 kg and varied more within (average of annual mass SD = 0.54 kg) than across cohorts (SD of annual average mass = 0.33 kg, Fig. S8). At the individual level, our multi-state capture-recapture modeling showed that fledging mass was positively related to the probability of surviving the juvenile phase, with equal support for linear and quadratic relationships (Table 1, models 16 and 17, LRT: both p < 0.001). The quadratic model predicts that juvenile survival plateaus at ca. 4 kg body mass (Fig. 4), and this relationship did not differ among years (Table 1). At the population level however, inter annual variations in juvenile survival were poorly explained by the average mass of the cohort (mean annual fledging mass, MFM, Table 1, model 35, %DEV = 7.2, slope ± 95%CI = 0.06 (0.04, 0.07)).

**Fig. 4.**
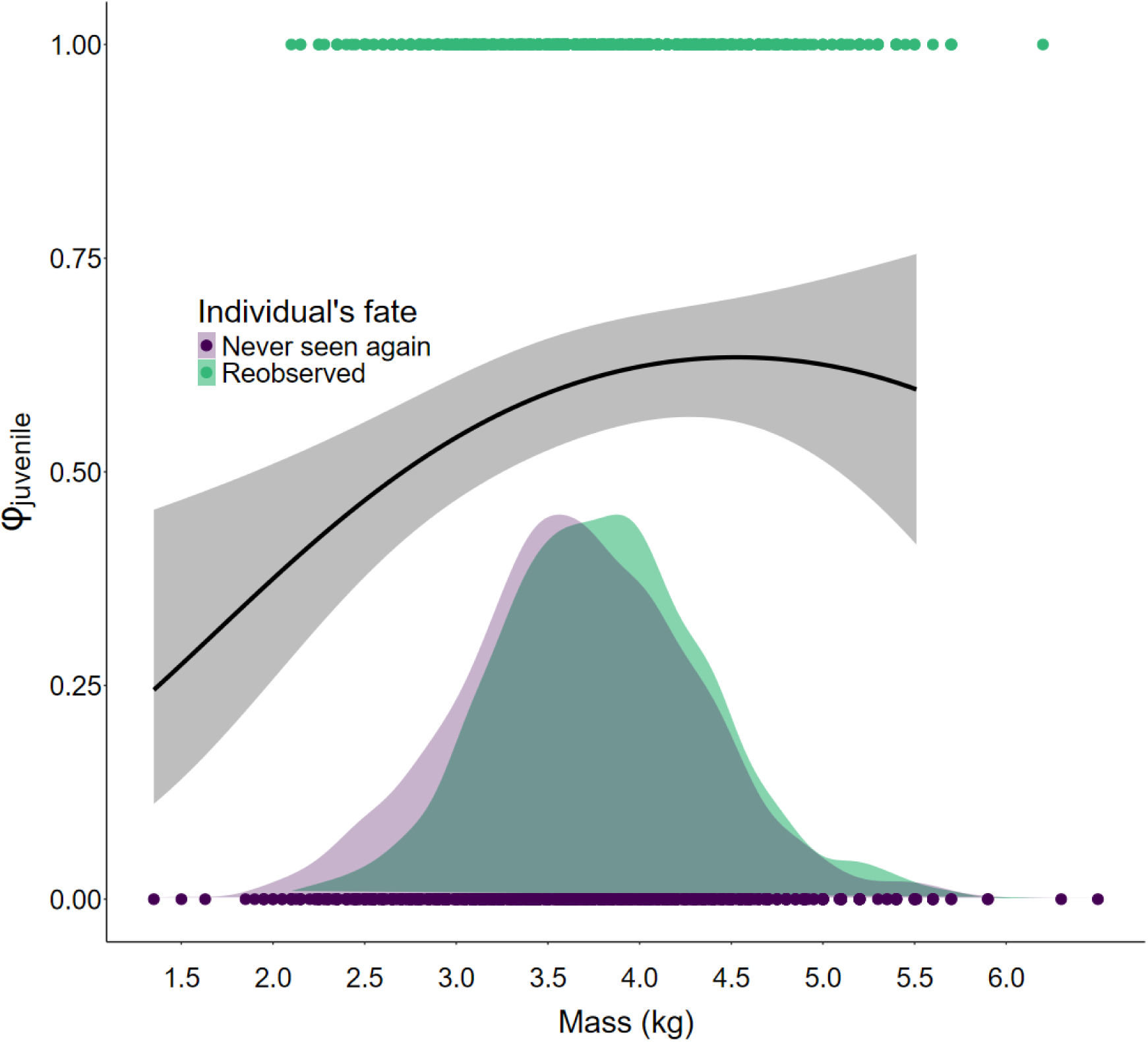
Model-predicted relationship between Adélie penguin juvenile survival (ɸ) and individual body mass at tagging. Modeling was carried out using the program MARK^98^. The predicted relationship is drawn from model 17 (Table 1). The density probability for each fate (reobserved vs. never seen again) is superimposed (shaded areas) as well as individuals (green and violet dots).

## Discussion

Using 17 years of individual electronic monitoring (2007-2023), we observed a loss of 2.5% per year in Adélie penguin juvenile survival over 14 years (2007-2020) in an Adélie Land population. This decline echoes those already observed in the only two other populations for which juvenile survival time series are also available (in Mac. Robertson Land, East Antarctica, -1.8% per year^31^, and in the Western Antarctic Peninsula, -1.3% per year^33^). By quantifying the survival of juvenile Adélie penguins for the first time in a region located thousands of kilometers away from other monitored colonies (3,400 km and 6,300 km, respectively), our results point to emerging negative trends across the continent. We also highlight potential mechanisms at play, since the decline in juvenile survival probability recorded in Adélie Land was associated with decreasing sea ice concentrations in the vicinity of the natal colony within two months after fledging. As sea ice declines are predicted to shrink Adélie penguin populations in the latter part of the century^17,18^, steady declines in juvenile survival may already foreshadow such shifts.

### Range-wide declines in juvenile survival

Hinke et al.^31^ were the first to report an annual decline in juvenile survival probability, with a decline of -1.3% per year between 1982 and 2000 for Admiralty Bay (King George Island, Western Antarctic Peninsula). Survival probability at this site then seemed to stabilize up to 2011, although a change in marking methodology (from aluminum to stainless steel flipper bands) and high inter-annual variability in survival probabilities made it difficult to compare the two periods^31,32^. More recently, Emmerson and Southwell^33^ extended the analysis of the capture-recapture dataset from Béchervaise Island (Mac. Robertson Land, East Antarctica^36^) to reveal that juvenile survival had also declined (-1.8% per year, 1992-2015) in this population located halfway between the Antarctic Peninsula and Adélie Land (Fig. 2).

With an annual decrease of -2.5% between 2007 and 2020 in Adélie Land, our study adds to the picture of declining juvenile survival across the species range (Fig. 2). Juvenile survival has now declined at all three locations where time trends have been investigated (Fig 2). The time series used here (14 years) is shorter than those for Admiralty Bay (18 years) and Béchervaise Island (24 years), but it is worth noting that the -2.5% annual decline at Pointe Géologie exceeds the average inter-annual variability in survival probabilities (1.79%) as in Béchervaise Island (1.80% vs. 1.57%) but unlike Admiralty Bay (1.30% vs. 2.90%).

Permanent dispersion of marked individuals away from study sites can bias the results of capture-recapture studies when incorrectly accounted for^41^. Here, our deployment of a mobile RFID antenna network, radiating 10 kilometers around the main study colony (Fig. 1), allowed a robust estimation of survival probability despite local dispersion. Long-distance permanent dispersion of juveniles, which could bias our survival estimates low, remains nonetheless largely unknown and should be addressed in the future.

### Drivers of juvenile survival

The existence of a relationship between Adélie penguin population dynamics and sea ice is well established^13,18,40,42^. However, despite the general importance of juvenile survival in driving population trends^33,43,44^, previous studies have found relatively limited evidence for a relationship between SIC and juvenile survival^31,33,36^. Here, unlike for the two populations previously studied^31,33^, we found that local sea ice concentration within 200 km of the natal colony in the timeframe of one to two months after fledging (April) accounted for up to 71% of the temporal variation in survival probability (Fig. 3). Specifically, higher SIC in April had a positive effect on juvenile survival. The timing of this relationship is consistent with a survival bottleneck occurring in the weeks following fledging, as identified for Adélie and other *Pygoscelis* penguins in the Antarctic Peninsula^45^. A mortality peak immediately following fledging is also consistent with previous studies in mammals^46,47^ and birds^48,49^.

Our finding that SIC in April, but not in March, was correlated to juvenile survival may be particularly informative on the mechanisms linking sea ice to the survival of young Adélie penguins. Although the winter freeze-up of the Antarctic Ocean begins in March, sea ice usually consolidates markedly and expands equatorward in April^50,51^ (see also Fig. S10). At the onset of winter, newly fledged Adélie penguins could then rely on such sea ice formation, especially dense landfast ice (Fig. 3, Fig. S11). By decomposing April SIC into its loose (the marginal ice zone, between 15 and 80% SIC) and dense (80-100%) fractions, we were able to show that juveniles survived better in years when sea ice was made of large, fully frozen areas, rather than patchy ice (Fig. 3B, 3C, Fig. S11).

Although only biologging studies may be able to determine how juvenile penguins exploit sea ice features, we hypothesize that dense sea ice could provide more suitable resting grounds for fledglings, in a context where they are at high risk of energy reserves depletion^45^. Furthermore, extensive fast ice areas contribute to the formation of polynyas^52^, where primary productivity is highest at the end of the summer^53^ and could thus concentrate penguin’s prey. Large amounts of loose sea ice may therefore impair the detection and exploitation of food -abundant areas by inexperienced penguins, likely to display lower foraging abilities compared to adults as shown in king penguins (*Aptenodytes patagonicus*)^54^. Finally, patchier sea ice may favor predation, especially by leopard seals (*Hydrurga leptonyx*)^55^, although predation pressure during fall and winter has never been investigated. During the winter, adult Adélie penguins tend to be associated with dense sea ice areas^56^, with higher SIC promoting higher survival to the next breeding season^37^. For juveniles, however, only two studies so far have investigated their post-fledging movements^45,57^, while their diving behavior and association with specific sea ice types remain completely unexplored.

While the major challenge for juvenile Adélie penguins may occur soon after fledging^45^, we also found that colder windchill temperatures during their first winter at sea (April-September) were associated with lower survival rates. A likely explanation involves the progressive build - up of thermoregulatory abilities that can continue up to several months into the juvenile (post-fledging) phase^58^. Although we cannot exclude that indirect rather than direct effects are linking winter windchill temperatures to juvenile survival, our results corroborate previous findings from another colony located several thousands of kilometers away from Adélie Land ^33^. More spatial coverage may be needed before confirming the generality of this driver, but weather conditions during winter (as proxied by windchill temperatures), could emerge as an important predictor of juvenile survival in this species.

Body mass at fledging is a common predictor for juvenile survival in birds^61^, including penguins^62–64^. Here, we found that survival probabilities increased steadily with chick mass, up to a plateau at ca. 4 kg body mass. While this echoes previous findings for the species^62,65^, this is demonstrated for the first time in a setup unbiased by flipper banding^66^ and accounting for imperfect detection. Importantly, the benefits of fledging at a higher mass were similar across years, suggesting that this advantage may be maintained across a range of environmental conditions (see also^62)^. While in the case of Pointe Géologie post-fledging sea ice conditions effects clearly superseded those of fledging mass in explaining interannual variations in juvenile survival, fledging mass could connect changes at other temporal scales to survival variations. For instance, both the diet^67,68^ and the foraging efficiency of breeders^69,70^ have been linked to sea ice conditions during the summer season or the previous winter, and these parameters may directly affect chick fledging mass^60,62^. Similarly, land-based, food-independent factors such as heavier precipitation and stronger winds in summer negatively affect chick fledging mass^59^, likely because of higher thermoregulatory costs^60^. In the Western Antarctic Peninsula region, where sea ice loss is associated with a poorer chick-diet^60^ and is stronger than in East Antarctica or in the Ross Sea^71^, these factors could act independently or combine with post-fledging sea ice conditions to reduce juvenile survival, thereby negatively affecting population trends^72^.

### Demographic consequences and perspectives

In long-lived species, the survival of young age classes can have major impacts on population dynamics^73^. For Adélie penguins, this is exemplified by declines in abundance of breeders that followed declines in juvenile survival probabilities in two distinct populations from the Western Antarctic Peninsula^31,32^ and East Antarctica^33^. As in the present study, it was not possible to identify the onset of the decline in juvenile survival in these two populations, since trends were observable right from the beginning of the monitoring (Fig. 2). Nonetheless, evidence from East Antarctica suggests that declining juvenile survival may not cascade into an abundance decline immediately, with a ∼15 years lag between the start of the juvenile survival time series and the breakup in the abundance time series^33^ (Fig. 2). Furthermore, events of low adult survival and repeatedly low breeding productivity may have acted in conjunction to generate the decline in abundance at Admiralty Bay^32^ and Béchervaise Island^33^, respectively. This may help explain why abundance at Pointe Géologie has not yet shown signs of decline^74^ (Fig. 2). However, given the occurrence of two massive breeding failures in the past decade alone^75^ and the stronger rate of decline in juvenile survival observed at Pointe Géologie compared to other studies and to local inter-annual variability in this vital rate, it is key to be closely monitoring the trends of the Pointe Géologie population in the coming years.

While Adélie penguin population declines for the Western Antarctic Peninsula are well documented^76–79^ and consistent with the stronger rate of warming in the region^80^, declines in other regions and notably in East Antarctica were not expected before the later part of the 21^st^ century^18^, with the Ross sea likely being the main refugia for the species^17^. However, our findings in the Western Pacific Ocean sector of Antarctica, combined with recent evidence that an entire metapopulation elsewhere in East Antarctica has already declined by almost half in the past decade alone^33^, suggest that changes could happen earlier than expected. Such rapid changes may challenge the adaptive abilities of the Adélie penguin. One of the main response mechanisms to these changes could be dispersal towards areas where environmental conditions remain or become favorable^18,81,82^. Although mid-range dispersion (< 120 km) is possible in Adélie penguins^83,84^, we still lack estimations of long-distance dispersion over time scales relevant with the current rate of environmental change.

In conclusion, our study has highlighted the emergence of similar negative trends in juvenile survival across the Adélie penguin range. While cross-population drivers behind these trends remain to be explored, recent range-wide studies^13,18,19^ provide a relevant template to achieve this goal. The next steps would first be to harmonize demographic databases across study sites and analyze them collectively. Specifically, quantifying the temporal variation in juvenile survival for the two Ross sea populations ^35,83,85^ is key. Completing the transition from flipper bands to RFID-tags^66^ could also facilitate comparisons of demographic rates among populations. Second, conducting biologging studies of juvenile movement and diving behavior, in conjunction with sea ice analyses (akin to^37^), is essential to uncover how inexperienced penguins exploit their environment. Third, rigorous efforts should be made to assess the dispersal capabilities of juvenile Adélie penguins under climate change, such as deploying autonomous RFID systems (Fig. S2) in populations adjacent to the main colonies where this method is used. With the forthcoming fifth International Polar Year (2032-2033) and the likely large-scale decline in the survival of juvenile Adélie penguins, a significant window of opportunity emerges for closely monitoring the demographic trends of vulnerable species and unraveling the underlying factors driving these trends.

## Methods

### Study area and design

Fieldwork was carried out at Pointe Géologie archipelago (Adélie Land, Antarctica; Fig. 1), close to the Dumont d’Urville research station (66°40′S, 140°01′E). This archipelago has hosted 30,000-50,000 breeding pairs of Adélie penguins annually for the last two decades^74^.

From the 2006-2007 breeding season onwards (breeding season will be referred to by the fledging year, e.g., 2007 in this case), Radio Frequency IDentification tags (RFID-tags) were implanted in chicks from a colony of about 270 breeding pairs located in a natural canyon (hereafter study colony). Unlike flipper bands^66,86,87^, RFID-tags are not known to negatively affect penguin vital rates. All chicks still alive right before fledging were tagged (n = 2,883 individuals). A few RFID-tagged chicks (n = 76) were found dead within the colony prior to fledging and were consequently excluded from subsequent analyses, bringing the final dataset to 2,807 individuals (range = 0-350/year, annual average = 187 from 2007 to 2021).

The study colony was fenced off in 2009 and instrumented with RFID gateways similar to those used in other penguin populations (Adélie penguins^88–90^; King penguins^91,92^; Southern rockhopper penguins^93^; Macaroni penguins^94^; Little penguins^95^). This setting made it impossible to miss tagged individuals visiting the colony. To account for dispersal from the study colony to neighboring colonies within a 10-km radius (Fig. 1), a grid of mobile RFID detection units (2-8 units, Fig. S2) was deployed every year from November to March, starting in 2013. Mobile RFID units were positioned at natural bottlenecks frequently used by penguins traveling between colonies and the sea. Units were moved every one to two weeks to maximize the number of detections. Among the 1061 individuals re-observed at least once after their tagging year, only 19 (1.8%) were detected exclusively by mobile RFID antennas, i.e. away from their natal colony. The mobile RFID-units resighting effort was more intensive closer to the study colony (< 200 m) but was performed up to 10 km from it (Fig. 1).

### Colony and individual-scale parameters

Colony-scale breeding productivity was defined as the number of fledglings, divided by the annual maximum number of adults inside the colony divided by two (a proxy for the number of breeding pairs). Individuals present in the colony were photo-counted weekly during the whole breeding season, starting in 2011. Cohort size was taken as the total number of RFID-tagged chicks that fledged. We ensured that observations from the study colony were representative of the larger-scale Pointe Géologie archipelago by comparing our local estimate of breeding productivity with data available from the literature for Pointe Géologie (2011-2017)^96^. Similarly, we ensured that our monitoring setup did not impact breeding abundance at the colony, by comparing the trend in number of breeding pairs in the study colony with that of two other undisturbed colonies located on the same island for the 2011-2024 period (Fig. S7).

All RFID-tagged chicks were weighed during tagging to the nearest 0.05 kg. The timing of tagging was adjusted to annual chick departure phenology (i.e. starting when the first molting chick was observed in the colony), ensuring mass was comparable across years. Each year, tagging occurred 10-15 days prior to fledging, making chick mass at tagging a reliable and consistent proxy for fledging mass^62^. In Pointe Géologie archipelago, fledging occurs from late February to early March.

### Capture Mark Recapture (CMR) modeling and statistical analyses

To model apparent survival and recapture probabilities of juvenile Adélie penguins, we constructed capture histories for all individuals (n = 2,807) fledged between 2007 and 2021 and subsequently observed between the 2009 and 2023 breeding seasons. Out of these 2,807 fledglings, 1,061 were observed again in subsequent years.

Given the dispersal of some birds from the study colony to neighboring colonies, and the higher resighting effort on and near the study colony, we implemented a multi-state framework to model survival, recapture, and transition probabilities^97^ using program MARK^98^ v.9.0. Different resighting probabilities among colonies were accounted for using a two-state classification: a local state (1) for the study colony and adjacent colonies within 200 meters, and a distant state (2) for all colonies located more than 200 meters from the study colony. This allowed closing the system and enhancing the robustness of survival estimates despite short-range dispersal dynamics. As the detection probability of 1-year-old individuals was near 0 (only a single bird was detected at age 1, Fig. S3), it was not possible to compute survival probability between ages 0 and 1. The first recapture occasion following tagging was thus deleted for all individuals (supplementary text). Consequently, juvenile survival was estimated between 0 and 2 years old.

We used U-CARE software^99^ to evaluate the goodness of fit (GOF) of the two-states time-dependent (JollyMoVe, JMV) model to our dataset^100^. The JMV model did not fit the original dataset because of transients (i.e. individuals never reencountered after tagging, Table S1a). We thus tested the GOF of the JMV model to a dataset where the first capture was set to 0 for each individual (i.e. effectively making capture history of each individual start at first reencounter following tagging, see supplementary text), thereby allowing to test the fit of a model with extended age-dependence on survival and recapture probabilities. The fit of the JMV model to this dataset was satisfactory (Table S1), albeit with indications of further age structure (i.e. > two age classes) in survival probabilities (significant test 3G.SM, Table S1b) and capture probabilities (almost significant Test M.ITEC, Table S1b). This validated the use of an initial umbrella model incorporating ≥ two age classes on recapture, survival, and transition probabilities (model 1, Table S3).

In Adélie penguins, presence at the breeding colony (and thus availability for detection) is conditional on age, with reproductively mature individuals more likely to visit colonies than pre-breeders. Survival is also age-dependent, with lower survival probability in juveniles compared to adults^35^. Earlier studies of known-age Adélie penguins have therefore accounted for age-structured recapture and survival probabilities, with either two^31^, three^36^, or five age classes^35^. Here, we started from a model with five age classes on survival, recapture, and transition probabilities, and explored all the possibilities in each submodel down to only two age classes. Finally, our starting (umbrella) model was adjusted to account for the low number of individuals detected in distant colonies. This model was framed with age variation only (no time effects) for all transition probabilities, and recapture probabilities in the distant colonies (state 2). The survival probabilities in both states (i.e. local and distant) were also set to be equal in that model, which appears as a reasonable assumption considering that all colonies belong to the same population of Pointe Géologie archipelago (Fig. 1). We successively looked for the most appropriate age and time structure for recapture, transition, and survival probabilities, using the previously retained best structure for each submodel. This selection process yielded an optimal model (model 13, Table S3) including two age classes (0-2 and 3+) and time effects within each age class for survival probabilities. Recapture probabilities were best modeled using three age classes in both states. The best structure included additional time effects in each of these three age classes for the local state only (Table S3). Transition probabilities were best modeled with state and age effects (five age classes in each state, Table S3). Because survival and recapture probabilities are not separable for the last year (2021) in time-dependent CMR models, this model allowed to estimate juvenile (0-2) survival from 2007 to 2020 only. Estimates of recapture probability in the local state were low and displayed high inter-annual variability for the first age class (mean ± SD: 0.33 ± 0.24, Fig. S9), but were high and stable in the last age class (mean ± SD: 0.96 ± 0.05, Fig. S9). Recapture probabilities in the distant state were estimated for the last age class only and were particularly low (0.02, SE = 0.009, Fig. S9). This model was then used as a basis for testing the effect of selected covariates on juvenile survival (see below).

### Covariate selection and definition

Candidate environmental variables were considered based on relevance to the Adélie penguin’s ecology and previous studies of Antarctic seabirds (Table S2). We first considered two large-scale indices, indicators of broad climatic variations in Antarctica and the surrounding Southern Ocean: the Southern Oscillation Index (SOI) and the Southern Annular Mode (SAM). Together, they affect wind patterns, temperatures, and sea ice dynamics, with complex regional-specific variability^7,101^. Such variability has been linked to opposite responses both among and within species. For instance, positive SOI values were previously linked to lower survival in adult Adélie penguins in Adélie Land ^34^ but no support for such an effect was found in Mac. Robertson Land^36^, the Western Ross Sea^35^, or the Antarctic Peninsula^31^. High SAM values were also found to affect the survival of other Antarctic seabirds, either positively (adult Cape petrel, *Daption capense*^102^; juvenile Emperor penguins, *Aptenodytes forsteri*^103^ or negatively (juvenile Snow petrels, *Pagodroma nivea*^104^). Monthly SOI data were downloaded from the NOAA National Centers for Environmental Information (https://www.cpc.ncep.noaa.gov/data/indices/soi) and daily SAM data from the NOAA Climate Prediction Center (https://www.cpc.ncep.noaa.gov/products/precip/CWlink/daily_ao_index/aao/aao.shtml).

Regional sea ice extent anomalies (SIEA) were also considered because of the quadratic relationship previously observed between SIEA and the survival of adult Adélie penguins in the Western Ross Sea^35^. Monthly data for the Western Pacific sector (90°E to 150°E), which encompasses our study site, were downloaded from the National Snow and Ice Data Center (NSDIC, https://nsidc.org/data/g02135/versions/3). To capture the conditions experienced by juveniles after fledging and during their first winter at sea, each index (SOI, SAM, SIEA) was averaged from March to September^36^, resulting in one value per year per index.

At the local scale, we considered both linear and quadratic effects of the average windchill temperatures (WCT) between April and September, based on the quadratic relationship previously observed between WCT and the survival of juvenile Adélie penguins^33^. Ten-day average data of minimum temperature, average wind-speed, and average relative humidity from the Dumont D’Urville weather station were downloaded from Meteo France (https://meteo.data.gouv.fr/) and used to calculate WCT following the Australian Bureau of Meteorology calculation (as in^33^).

Finally, we considered sea ice concentrations (SIC, average sea ice concentration in a specific area) at three different spatio-temporal scales (Fig. S4). As tracking data for juveniles is currently not available in our focal region, we considered broadly similar migration routes as those of adults previously studied at Pointe Géologie and found to gradually move westward after reproduction following the Antarctic coastal current^105^. This pattern is corroborated by the tracking of both juveniles and adults in another east Antarctica population^36,57^ and therefore likely in our study area. To reflect post-fledging (autumn) conditions experienced by birds near their natal colony, we considered March-April SIC between 135°E and 145°E (SIC.autumn). We also considered winter (May-July, SIC.winter) and late winter (August-September, SIC.latewinter) conditions, both between 110°E and 135°E. Sea ice concentrations were computed by averaging daily concentration values from 25×25 km grids downloaded from the NSDIC^106^ (https://nsidc.org/data/g02202/versions/4) over the spatio-temporal windows defined above. Following initial analyses, we further adjusted our sea ice metrics by first decomposing SIC.autumn into March and April SIC (SIC.march, SIC.april). Because different types of ice may affect seabirds differently^107^, we then calculated the April fraction of the total ice area (15-100% SIC) covered by the marginal ice zone (loose sea ice between 15-80% SIC, MIZFRAC.april) and the April fraction of the total ice area covered by dense pack ice (80-100% SIC, DENSEFRAC.april).

We also considered cohort size and average mass at tagging (as a proxy for fledging mass^62^) as likely intrinsic drivers of juvenile survival in Adélie penguins. Larger cohorts could reflect favourable growth conditions or be associated with lower predation pressure, as suggested in a previous study^33^, while fledging mass is a regular predictor for juvenile survival in birds^61^, including penguins^62–64^. The effect of fledging mass on inter-annual differences in survival probability was investigated by using mean fledging mass (MFM) for each cohort. Inter-individual, within-season effects of fledging mass were investigated, using chick mass as an individual covariate.

To avoid including covariates correlated with each other (multicollinearity)^25^, we checked for pair-wise correlations among our initial set of covariates (SOI, SAM, SIEA, SIC.autumn, SIC.winter, SIC.latewinter, WCT, MFM, cohort size). Only SIC.winter was correlated strongly (*r* > 0.7)^108^ with two other closely related sea ice metrics (SIEA and SIC.latewinter) so that we decided to drop this covariate from analyses. Remaining covariates were scaled to improve model convergence and facilitate parameters comparisons. We first investigated the effect of individual fledging mass on juvenile survival using Likelihood Ratio Tests (LRT), and included this parameter in further models if it improved model fit. We then tested annual covariates in isolation using ANODEV and considered them influential when the 95% confidence interval (CI) of the effect size did not overlap 0 and when they accounted for ≥ 20% of the temporal variation in survival rates^25^. The support for a linear trend in juvenile survival was assessed using ANODEV^25^ on a model set (i.e. constant, time, and trend models) that did not include fledging mass as an individual covariate in order to account for any contribution of fledging mass in driving potential temporal trends.

Homoscedasticity and normality assumptions were assessed for linear models. Values are reported as means ± SD unless mentioned otherwise. Statistical analyses other than the CMR modeling were conducted in R^109^ version 4.3.2. Timelines for the different datasets used in the study (RFID-tagging of chicks, resighting of RFID-tagged individuals, number of breeding pairs and breeding productivity data) are summarized in Table S4.

## Acknowledgements

We are deeply grateful to all the members of Project 137, including Benjamin Friess, Yvon Le Maho (former PI of 137-ECOPHY), Victor Planas-Bielsa, Claire Saraux, and all the wintering and summering field teams since the inception of this project in the field in 2005. We would also like to thank all the members of the missions in Dumont D’Urville since then, and the French Polar Institute-IPEV logistics team in Dumont d’Urville for their important and continuous support in the field.

## Author contribution

Author contributions follow the CRediT guidelines. In each section, authors are listed by order of appearance in the author list.

Conceptualization: TB, NL, CLB

Methodology: TB, RCH, AH, MBE, RCR, TR, NC, JC, MBU, NL, CLB

Software: TB, GB, RCH

Validation: TB, NL, CLB

Formal analysis: TB, GB, RCH, NL

Investigation: All authors

Resources: CLB

Data Curation: TB, GB, CCC, CLB

Writing – original draft preparation: TB, NL, CLB

Writing – review and editing: All authors

Visualization: TB

Supervision: NL, CLB

Project administration: CLB

Funding acquisition: DPZ, NL, CLB

## Data and material availability

Data and codes will be made available upon acceptance of the manuscript.

## SUPPLEMENTARY MATERIAL

## Supplementary Text

### Manipulating capture histories to account for near-zero recapture probability at age 1 and for Goodness-Of-Fit (GOF) testing

#### 1. Accounting for near-zero recapture probability at age 1

In capture-recapture modeling, the capture history (CH) of an individual is coded as a suite of numbers for each capture occasion. For example, if 0 = not seen; 1 = seen in state 1; and 2 = seen in state 2, the individual A presented in the table below was tagged in state 1 in year 2, not seen in year 3, seen in state 1 in year 4 and seen in state 2 in year 5.

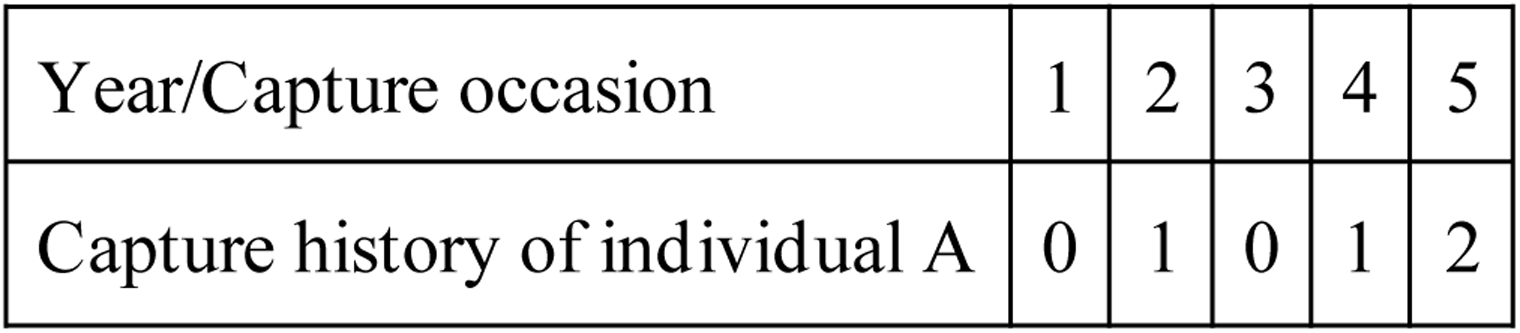

The CH of individual A can be written as: “01012”. Because recapture probabilities in the year following tagging (year 3 in the case of individual A) were near 0 in our study (only one individual was detected at one year old over our entire study period, Fig. S3), we rewrote CH of all individuals by removing the year after tagging. For example, the CH of individual A was transformed from “01012” into “0112”.

This allows accounting for the fact that with so few individuals seen again one year after fledging, it is not possible to estimate survival probability (ɸ) from age 0 to age 1 (ɸ 0→1). Instead, we estimated survival from age 0 to age 2 (ɸ 0→2). This is the typical “juvenile survival” in capture-recapture studies where individuals are not available for detection before two years after fledging. Under the specific (but untestable in this case) hypothesis that ɸ 0→1 = ɸ 1→2, survival from age 0 to age 1 (ɸ 0→1) can nonetheless be estimated as sqrt(ɸ 0→2).

#### 2. GOF testing

There is no direct GOF test for age-dependent, multistate capture-recapture models^99^. However, the presence of transients (individuals never seen again after tagging) is a common source of age-dependence in recapture probabilities. To quantify how much of the lack of fit of a time-dependent (JollyMoVe, JMV) model can be attributed to transients, a common strategy is to test the GOF of the JMV model to a dataset where transients are removed by replacing the first capture of each individual by 0 in their capture histories. For example, in the dataset where transients are removed, the CH of individual A would be “00012”.

When the GOF of the dataset without transients is satisfactory, the original dataset (i.e. including transients) can be used but recapture probabilities must be specified as age-dependent in the starting model, therefore accounting for transience.

**Fig. S1.**
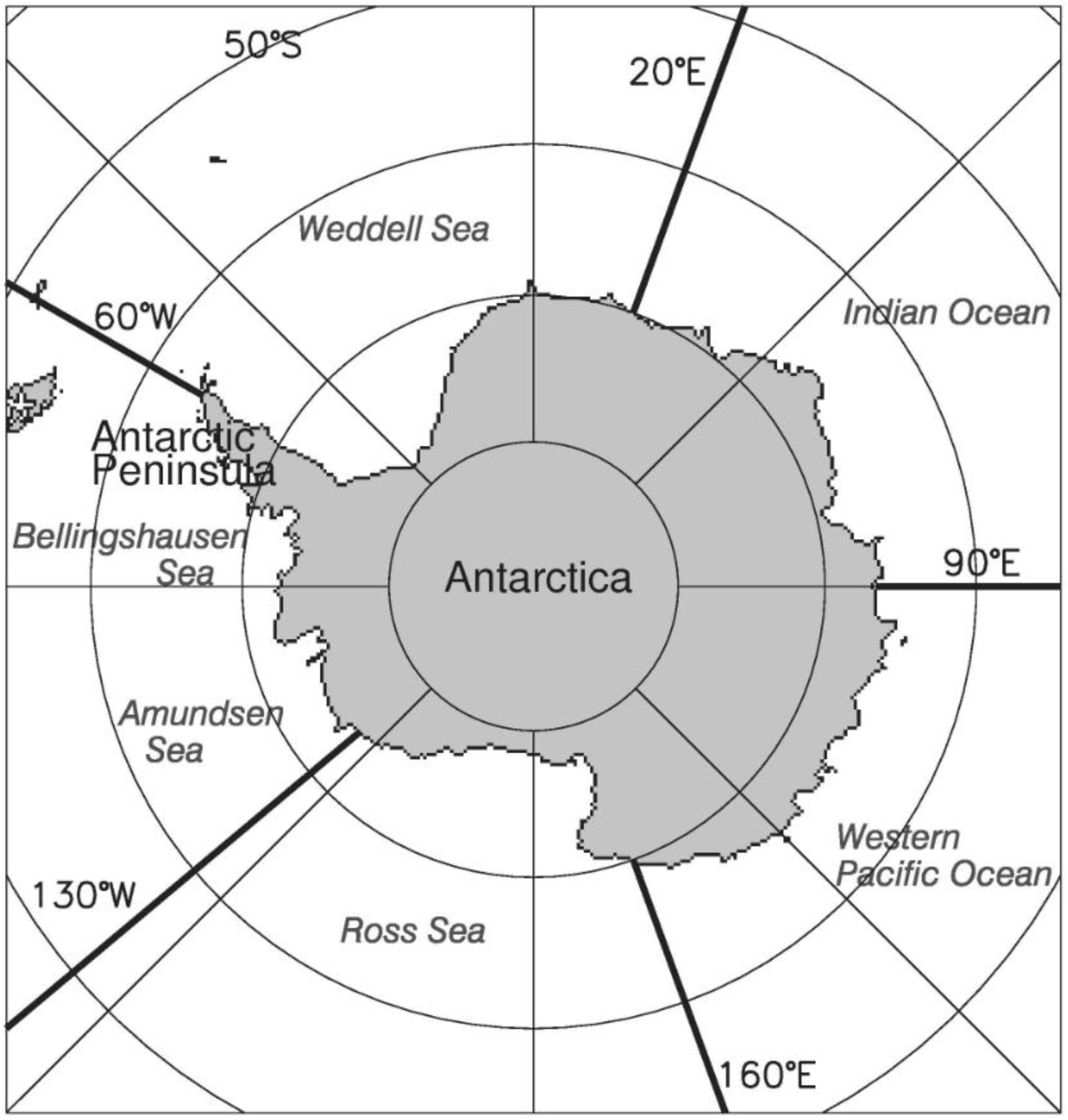
Sectors of Antarctica used for large-scale sea ice analyses. Figure extracted from^71^.

**Fig. S2.**
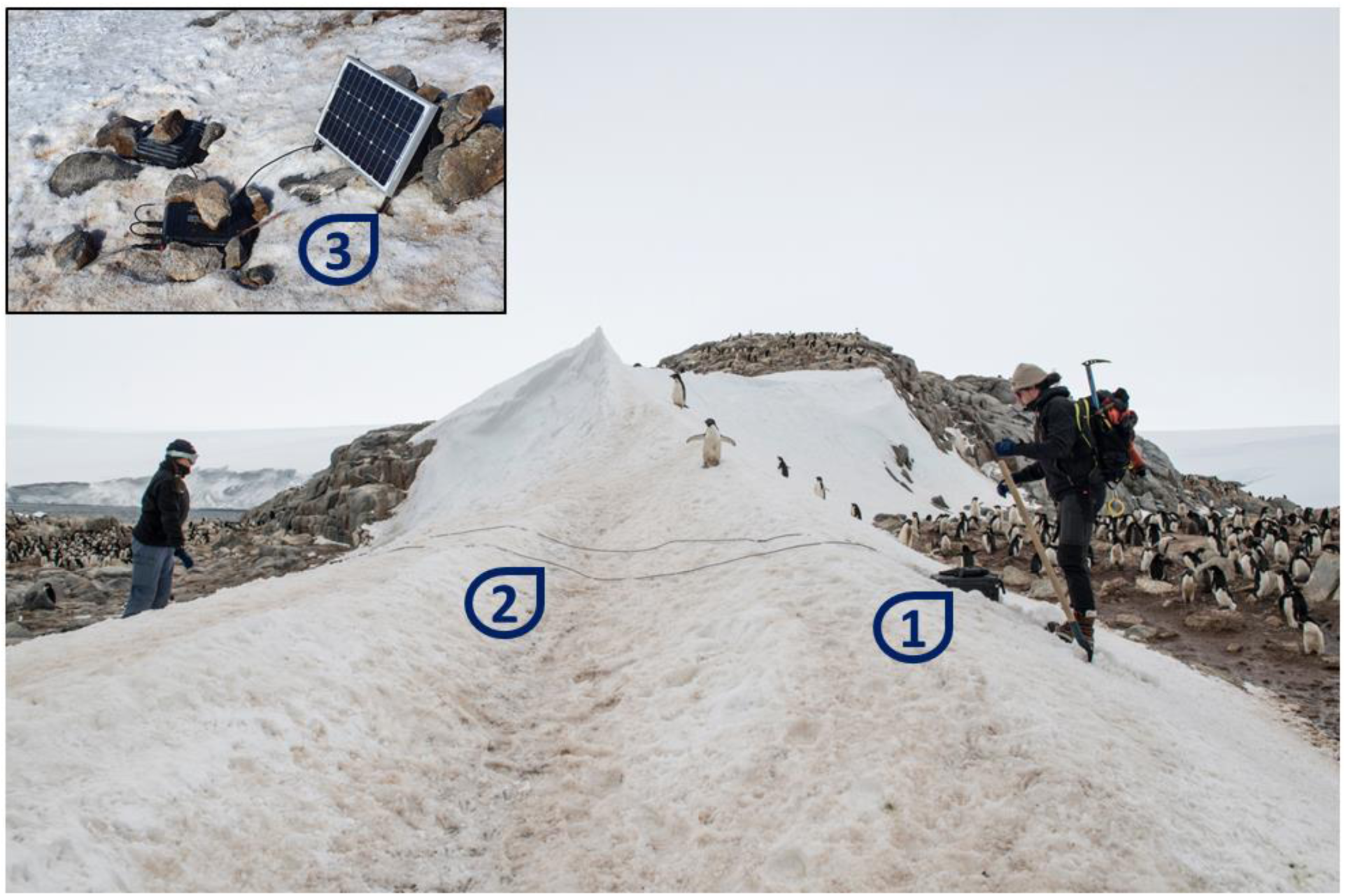
A mobile RFID detection unit being deployed on an Adélie penguin (*Pygoscelis adeliae*) passageway near a breeding colony in Adélie Land. This system allows the passive detection of RFID-tagged individuals away from the main study site where tagging is conducted. Detection data and batteries are stored in the acquisition box (1), to which an antenna is affixed and buried in the snow on penguin passageways (2, burying in process in this picture). To increase running time up to several weeks during the summer season, the system can be fitted with solar panels (3). ⓒ Gregory Tran (main panel), Téo Barracho (inset), Institut Polaire Français - French Polar Institute

**Fig. S3.**
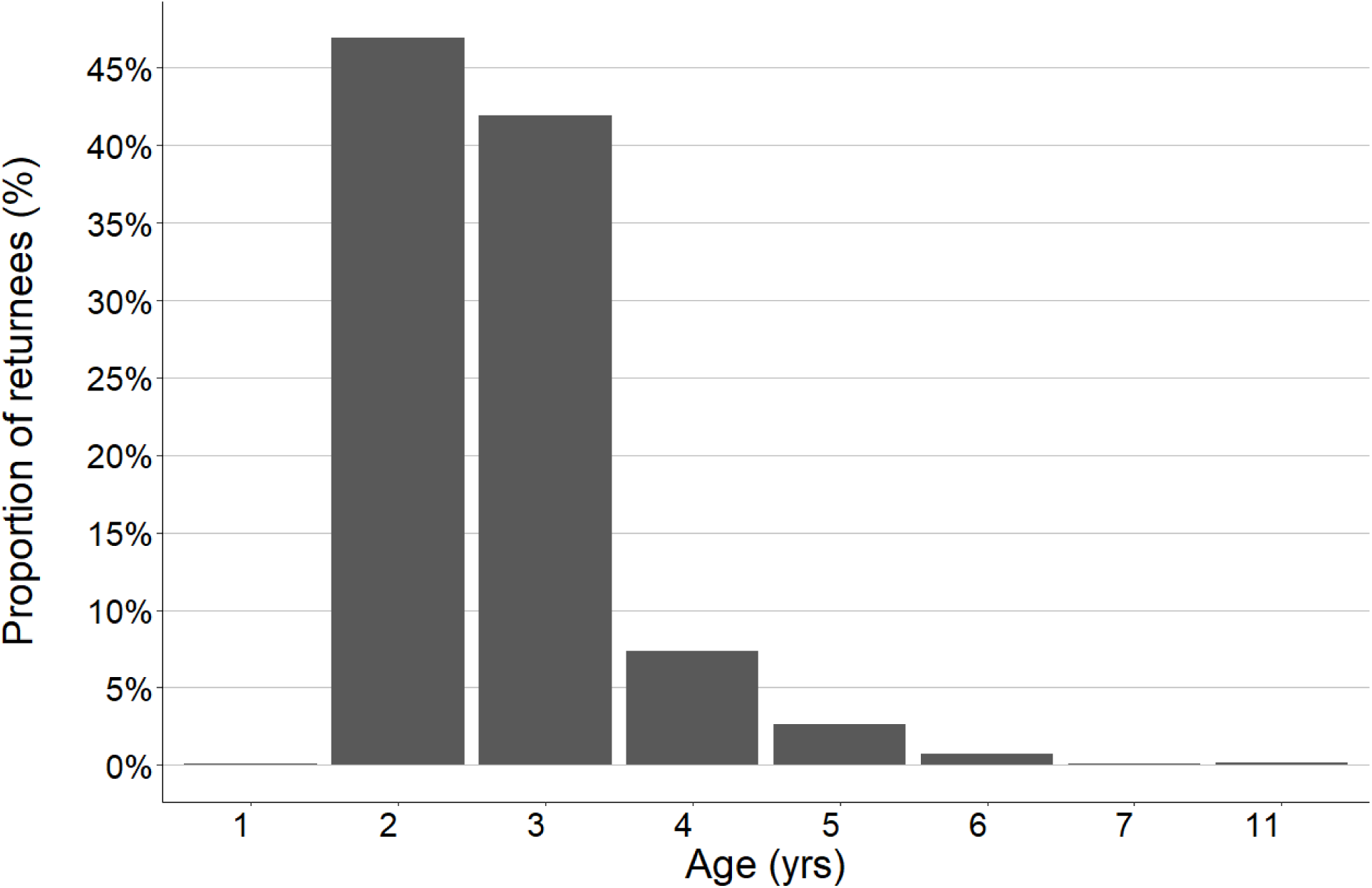
Age of first return at the colony after fledging for RFID-tagged Adélie penguins in Pointe Géologie archipelago, Adélie land, Antarctica (2007-2020)

**Fig. S4.**
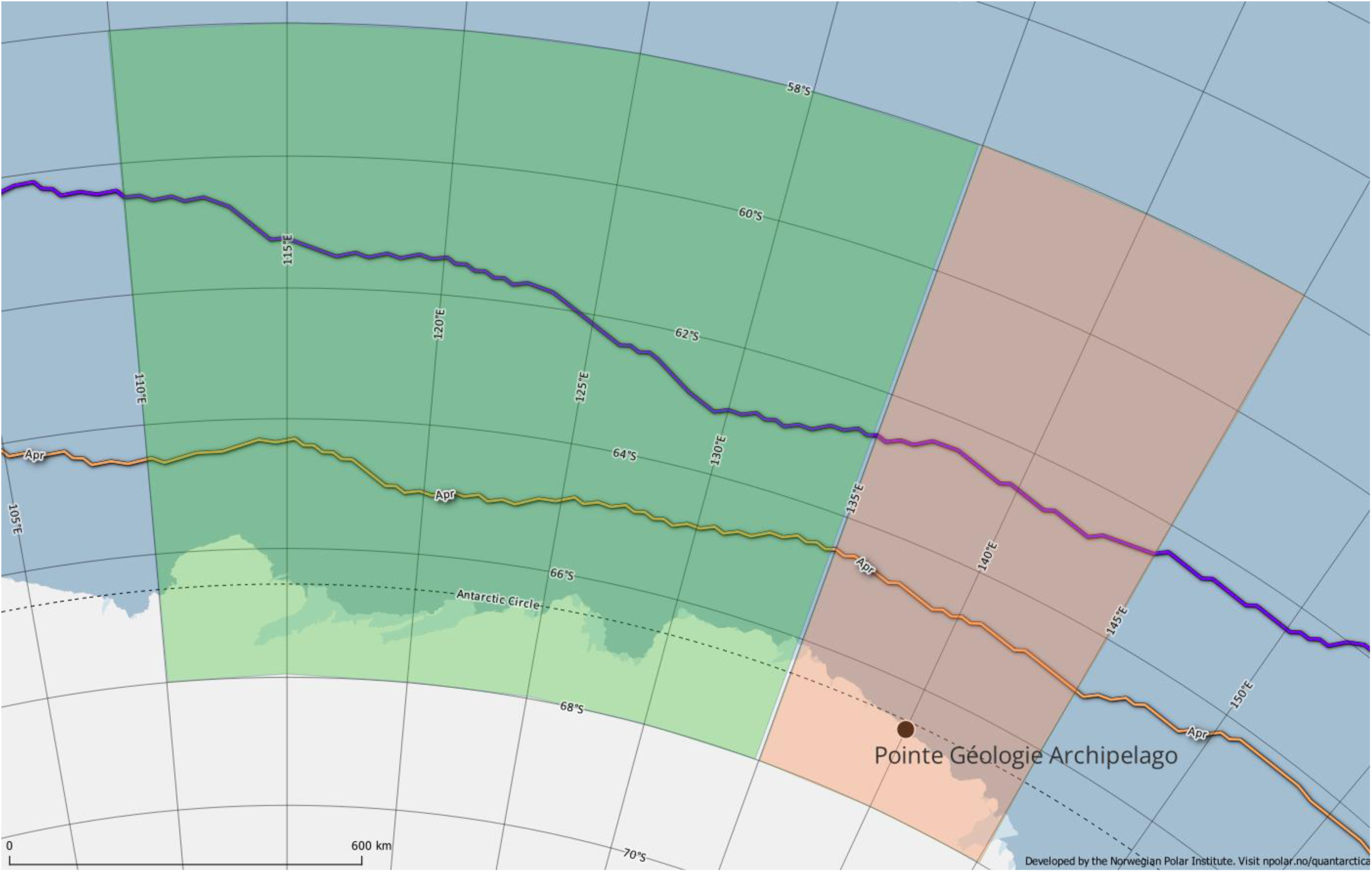
Spatial windows considered for estimating the sea ice concentrations covariates for autumn (orange), and winter/late winter (green). The yellow line depicts the average April sea ice extent and the blue line the average maximal sea ice extent (both over the 1981-2010 period). Map produced using Quantarctica^110^(https://www.npolar.no/quantarctica/)

**Fig. S5.**
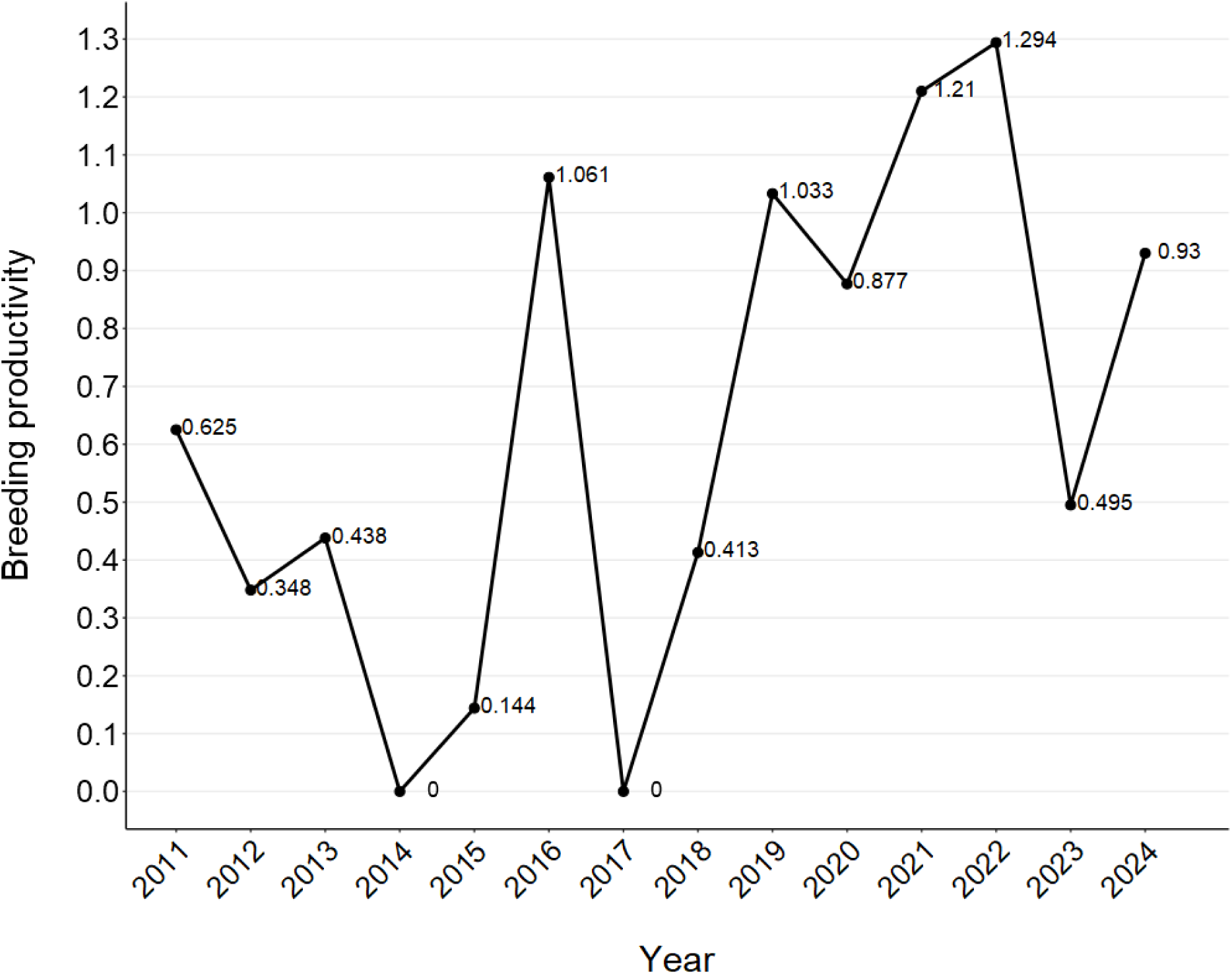
Time series of breeding productivity for the study colony (2011-2024, average ± SD = 0.63 ± 0.44 chicks per breeding pair). Breeding productivity was calculated as the number of chicks fledged per breeding pair. The number of breeding pairs was proxied as the annual maximum number of adults inside the colony (from weekly photo counts), divided by two. Breeding productivity may be superior to 1 in some years because Adélie penguins can raise two chicks^40^.

**Fig. S6.**
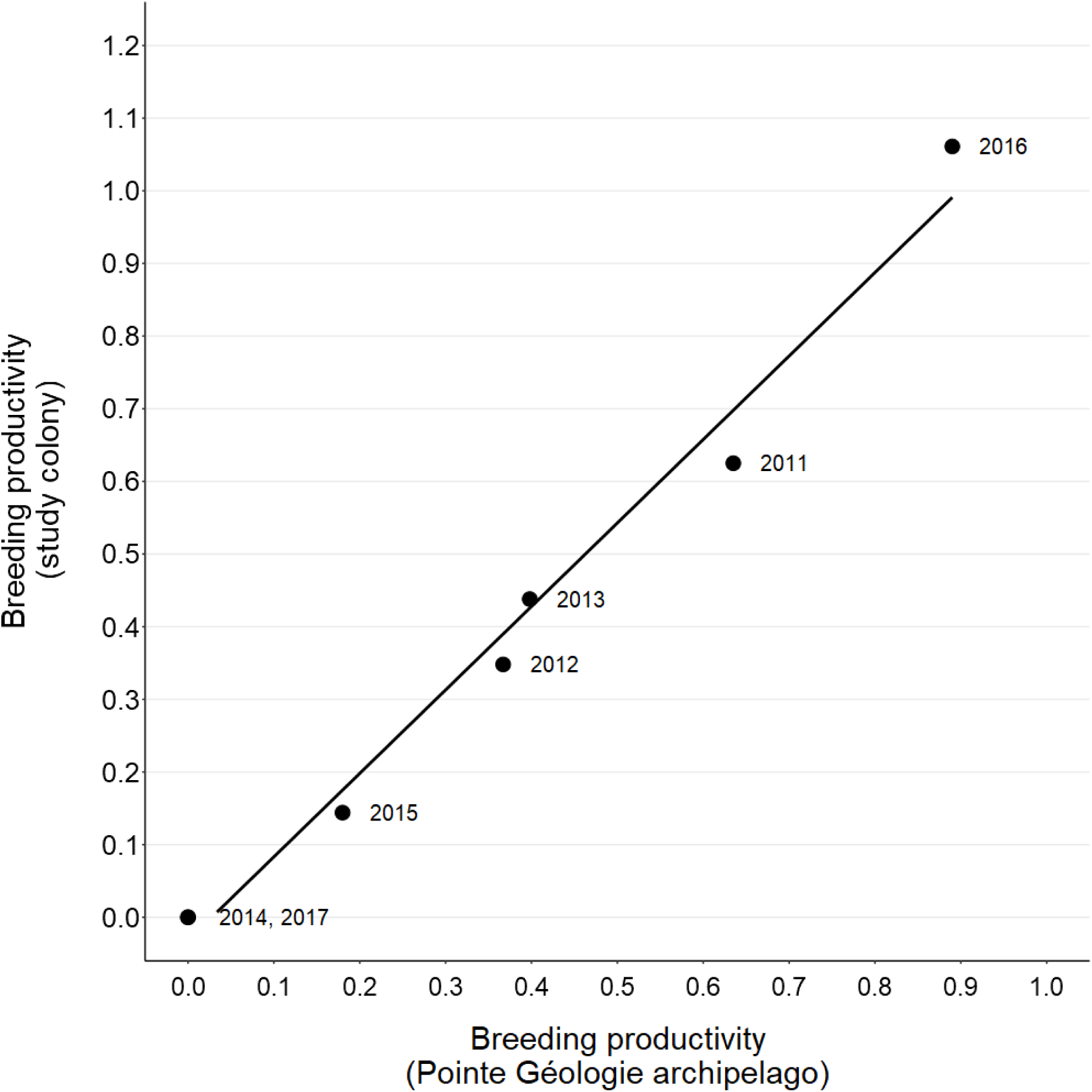
The annual breeding productivity (chicks per pair) of the study colony correlates with that of the whole population of Pointe Géologie archipelago, Adélie Land, Antarctica (2011-2017, Pearson correlation: r = 0.99, p < 0.001). Data for Pointe Géologie was extracted from^96^. Breeding productivity may be superior to 1 in some years because Adélie penguins can raise two chicks^40^.

**Fig. S7.**
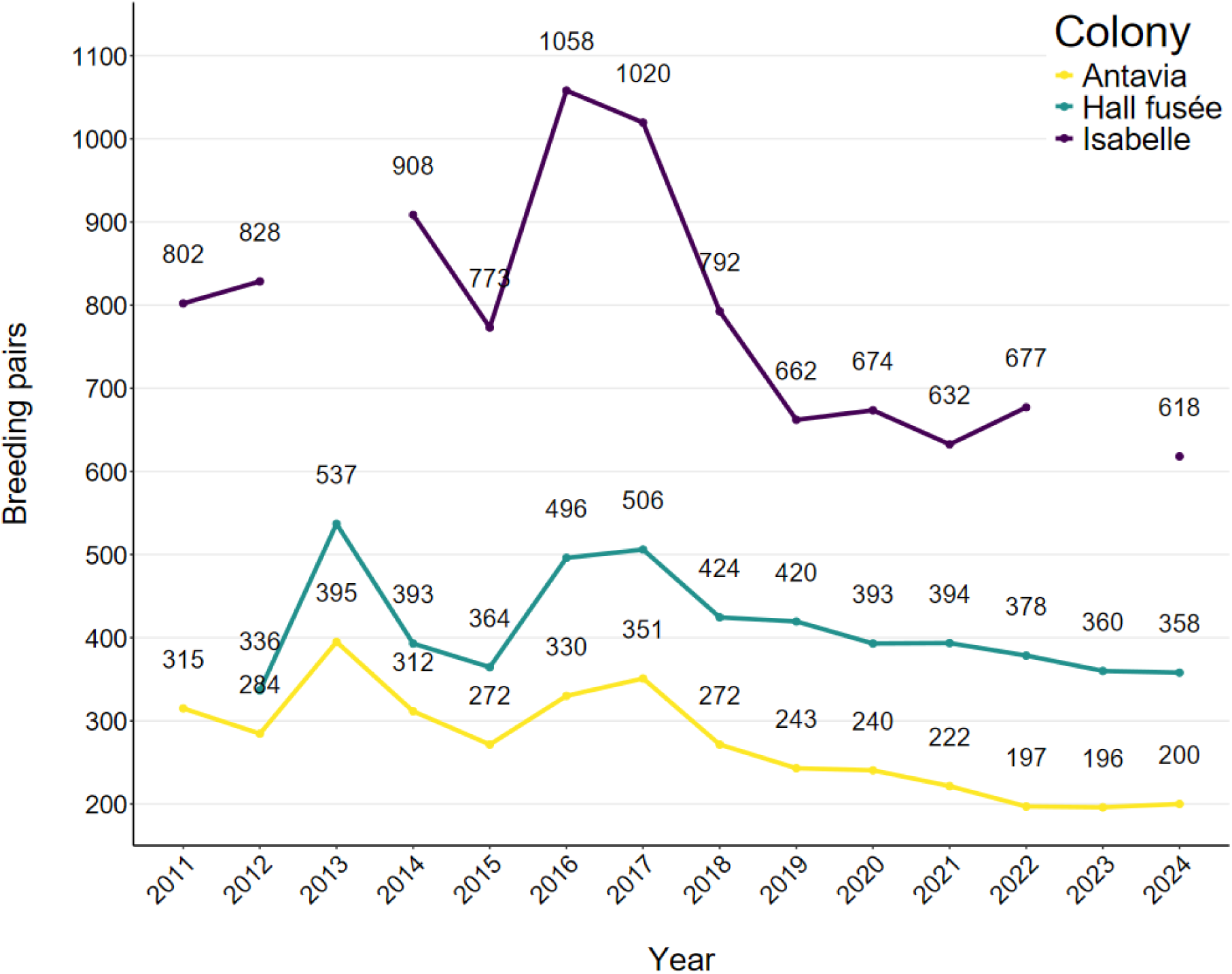
Number of breeding pairs (as proxied by the annual maximal number of adults counted during weekly surveys) in three neighboring Adélie penguin colonies in Pointe Géologie archipelago, Adélie Land, Antarctica. These three colonies include the study colony (Antavia) and two control colonies located on the same island (Hall Fusée and Isabelle) where no monitoring activities are conducted except for weekly photo counts. The number of breeding pairs in the study colony (Antavia) averaged 273 ± 62 between 2011 and 2024, and interannual variation remained comparable with that of the two control colonies (Spearman rank correlations, both p < 0.022).

**Fig. S8.**
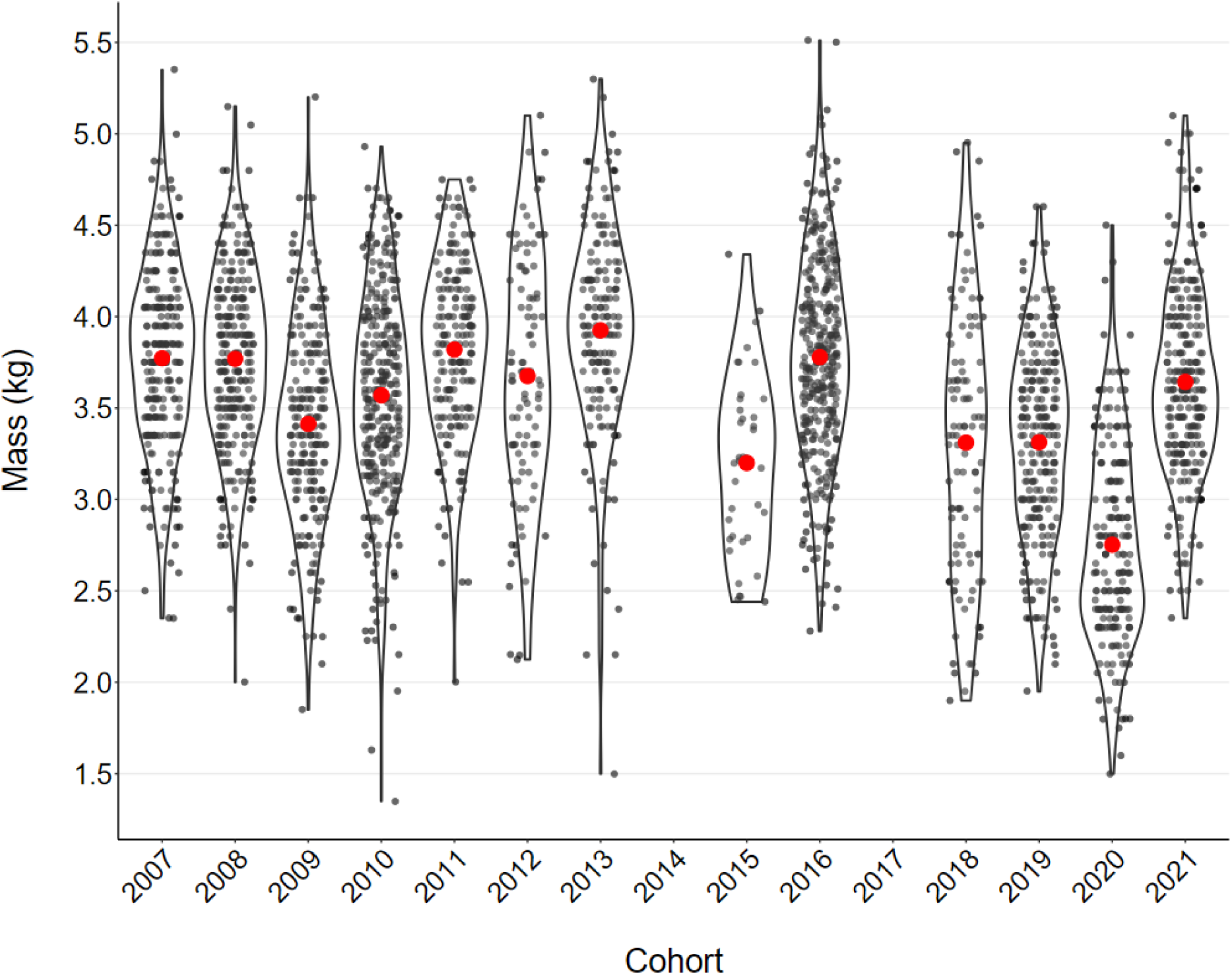
Chick body mass at the time of RFID-tagging across cohorts (2007-2021) for Adélie penguins born in the study colony (Pointe Géologie archipelago, Adélie Land). Red dots represent annual means. Tagging consistently occurred 10-15 days prior to fledging, thus making mass at tagging a reliable proxy for fledging mass^62^.

**Fig. S9.**
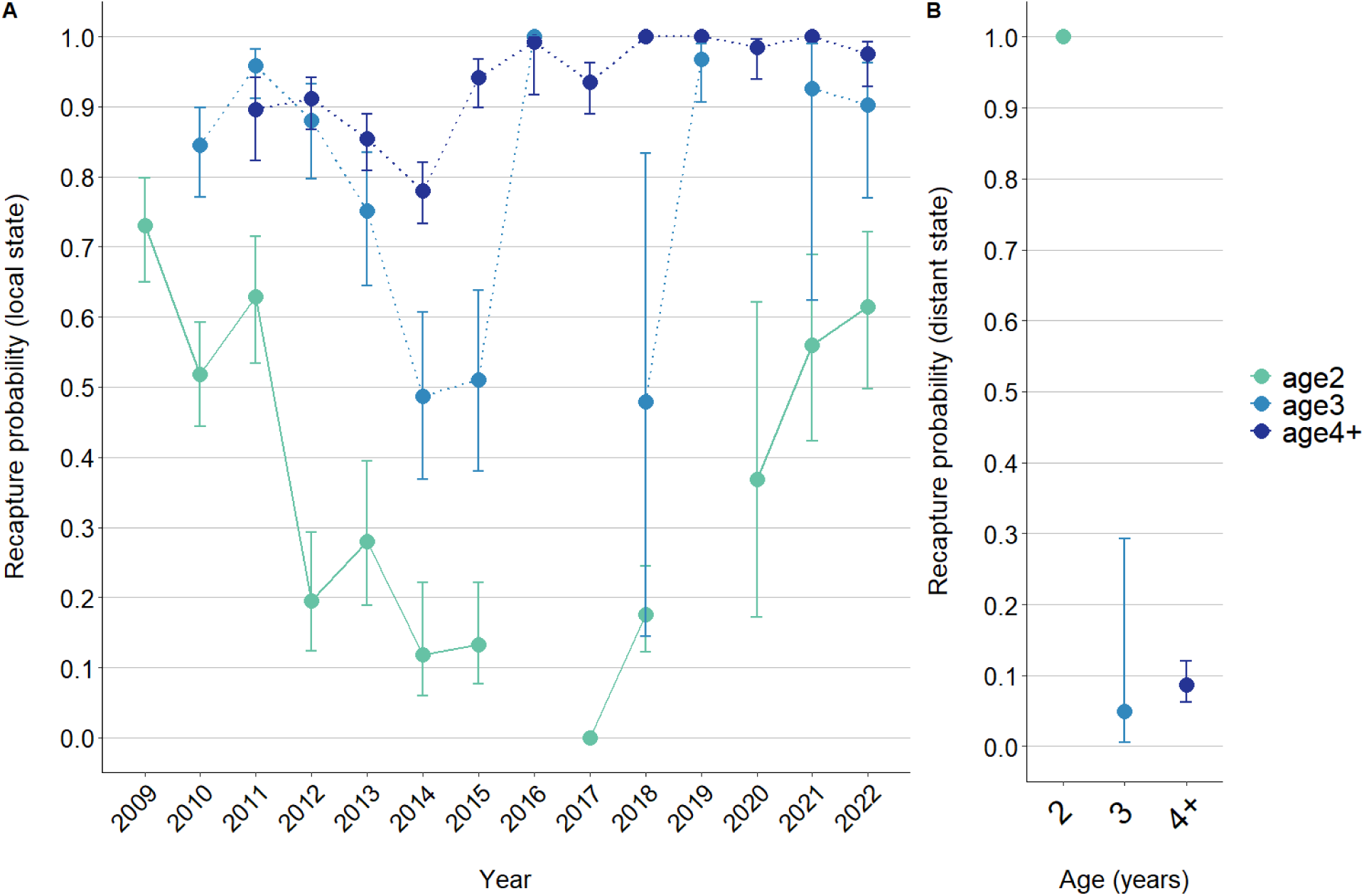
Estimated recapture probabilities of known-age RFID-tagged Adélie penguins in the local (A) and distant (B) states. States refer to spatial locations of colonies where individuals are detected. The local state includes the study colony and adjacent colonies (< 200 m), while the distant state includes colonies elsewhere in Pointe Géologie archipelago (see Fig. 1 in the main text). Error bars indicate ± SE.

**Fig. S10.**
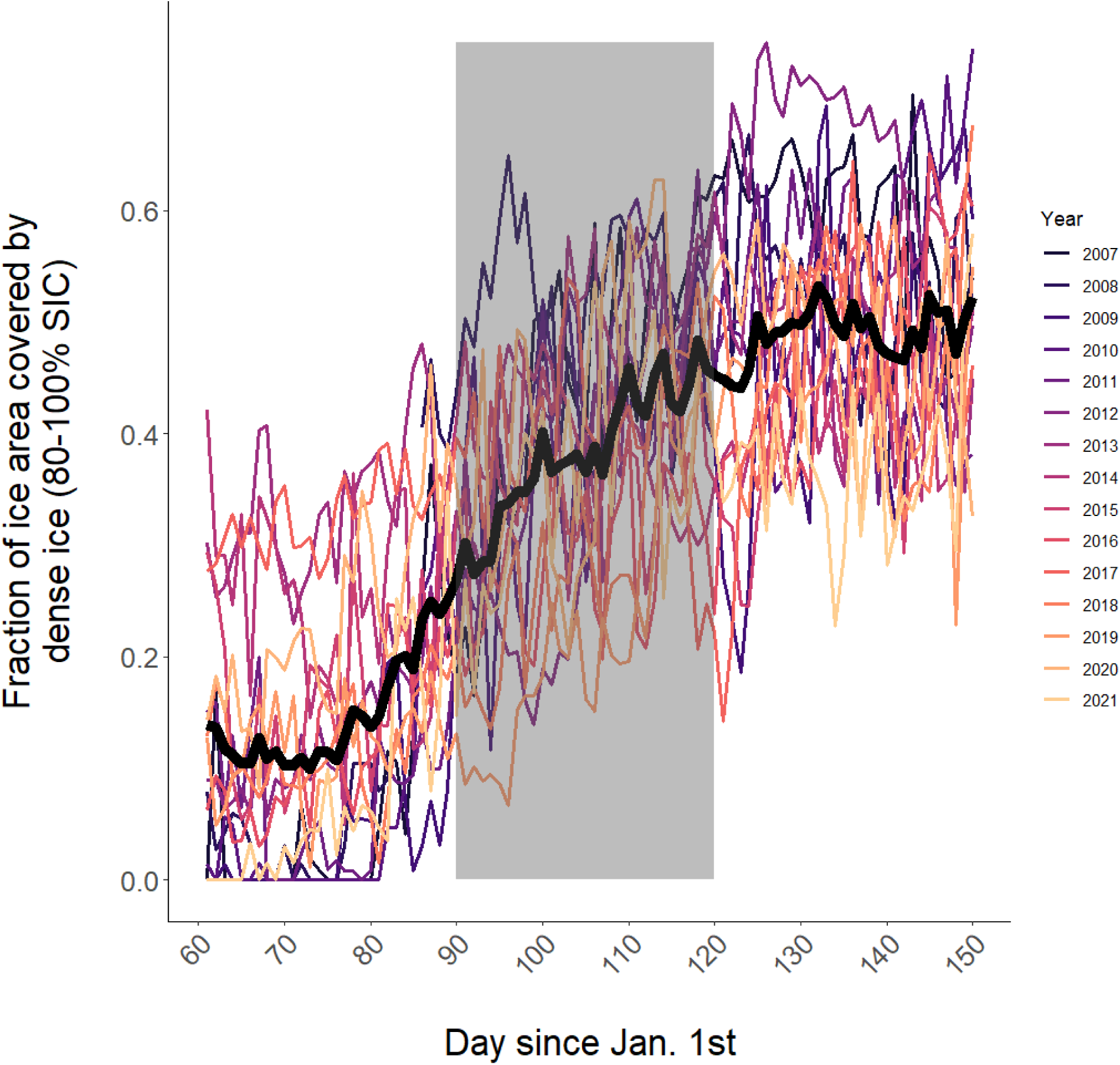
Annual increase in the proportion of dense sea ice (80-100% sea ice concentration) in the vicinity of Pointe Géologie archipelago (∼200 km) between March and May of each year (2007-2021). The gray shaded area represents the month of April, where the increase in dense ice fraction is strongest. Daily sea ice concentration data over 25×25 km grids was downloaded from the NSDIC^106^ (https://nsidc.org/data/g02202/versions/4).

**Fig. S11.**
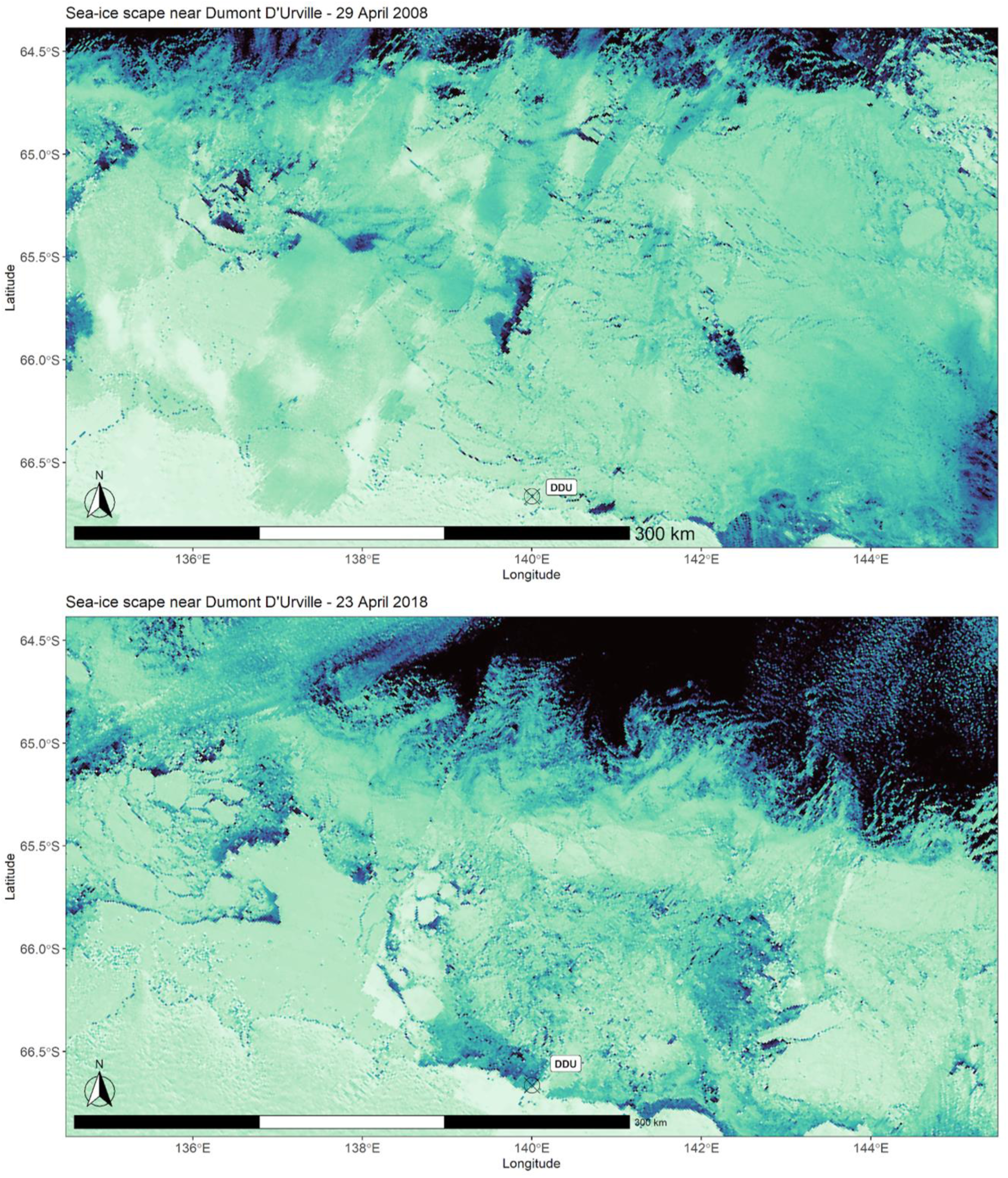
Sea ice-scape at the end of April near Pointe Géologie archipelago in two years of contrasting survival probabilities for juvenile Adélie penguins (2008, high survival probability; 2018, low survival probability. MODIS satellite images were downloaded from the NASA Worldview application (https://worldview.earthdata.nasa.gov), part of the NASA Earth Observing System Data and Information System (EOSDIS).

**Table S1:** Results of Goodness-of-fit tests for the multistate capture-recapture dataset used to estimate Adélie penguin juvenile survival probability at Pointe-Géologie archipelago, Adélie Land, Antarctica). df, Degrees of freedom; ĉ, Deviance inflation factor.

| Test | $\chi^2$ | *pval* | df | $\hat{c} = \chi^2/Df$ |
| --- | --- | --- | --- | --- |
| A) Original dataset |  |  |  |  |
| WBWA | 2.469 | 0.7 | 4 | 0.617 |
| 3G.SR | 1827.802 | <0.001 | 12 | 152.317 |
| 3G.Sm | 1393.189 | <0.001 | 35 | 39.805 |
| M.ITEC | 18.37 | 0.005 | 6 | 3.062 |
| M.LTEC | 6.831 | 0.2 | 5 | 1.366 |
| JMV model | 3248.661 | <0.001 | 62 | 52.398 |
| B) Dataset where 1st capt. set to 0 (no transients) |  |  |  |  |
| WBWA | 3.428 | 0.5 | 4 | 0.857 |
| 3G.SR | 14.744 | 0.324 | 13 | 1.134 |
| 3G.Sm | 47.702 | 0.0 | 31 | 1.539 |
| M.ITEC | 12.541 | 0.051 | 6 | 0.805 |
| M.LTEC | 6.015 | 0.3 | 5 | 0.919 |
| JMV model | 84.429 | 0.017 | 59 | 1.431 |

**Table S2.**
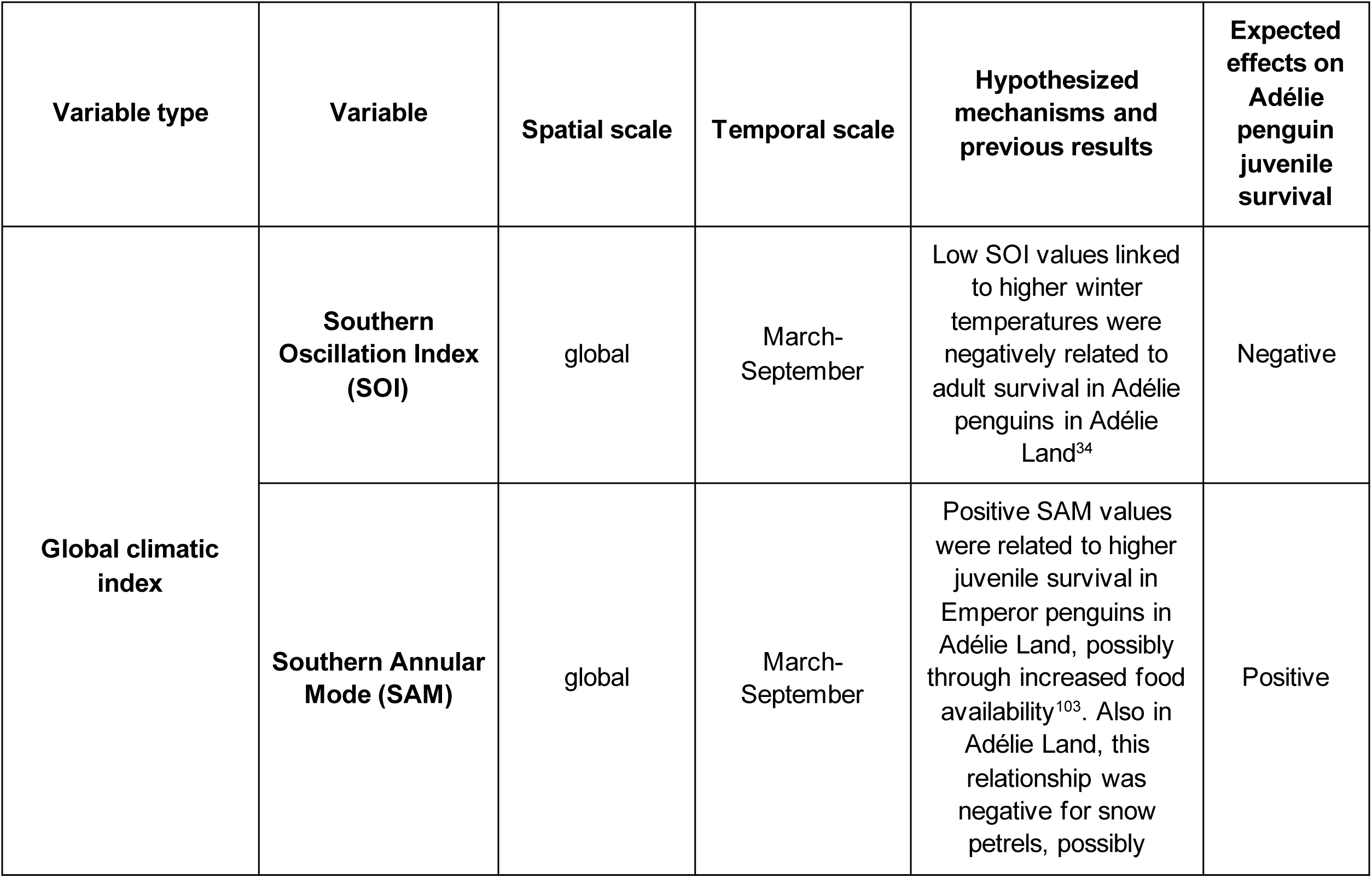
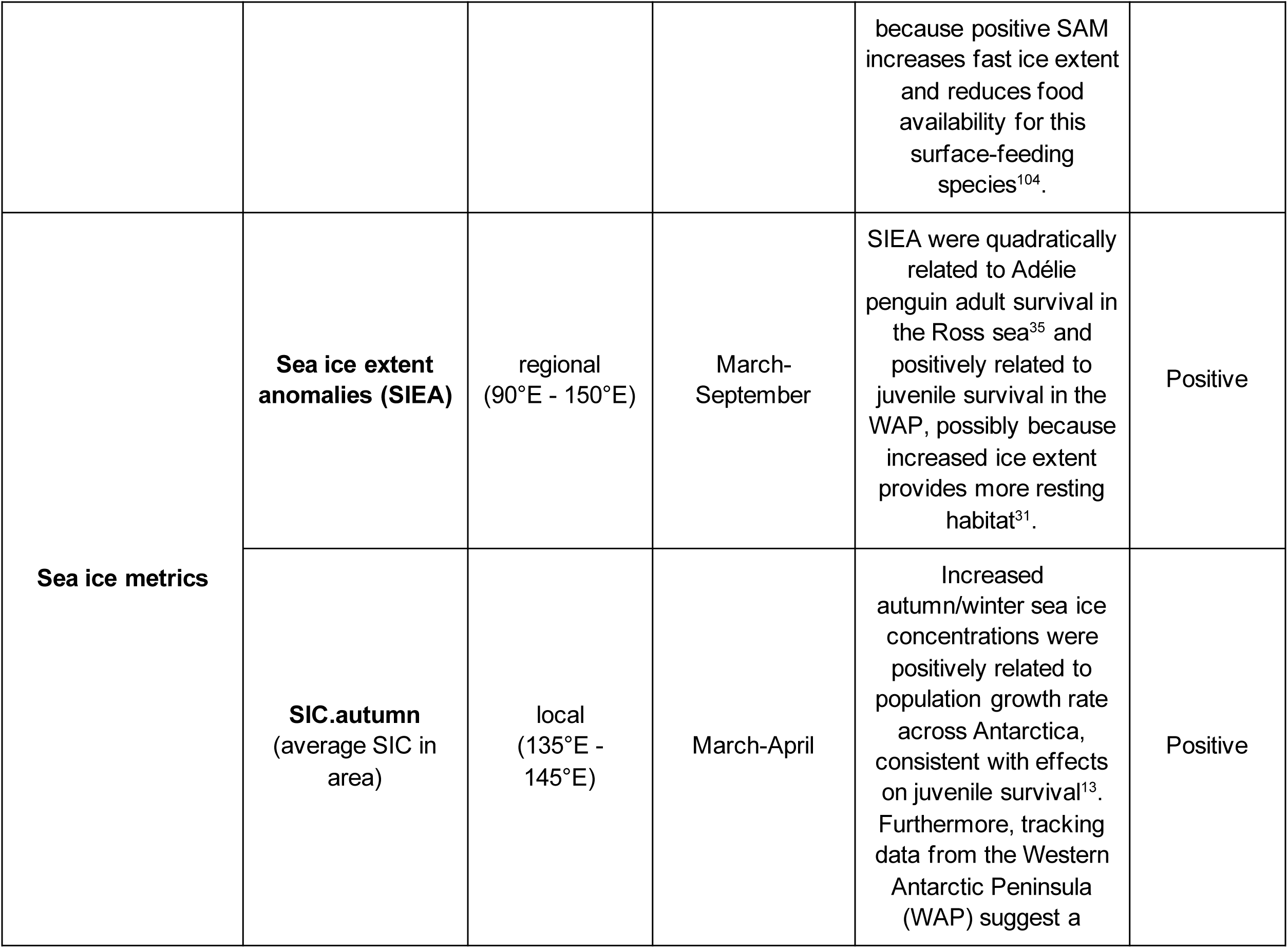
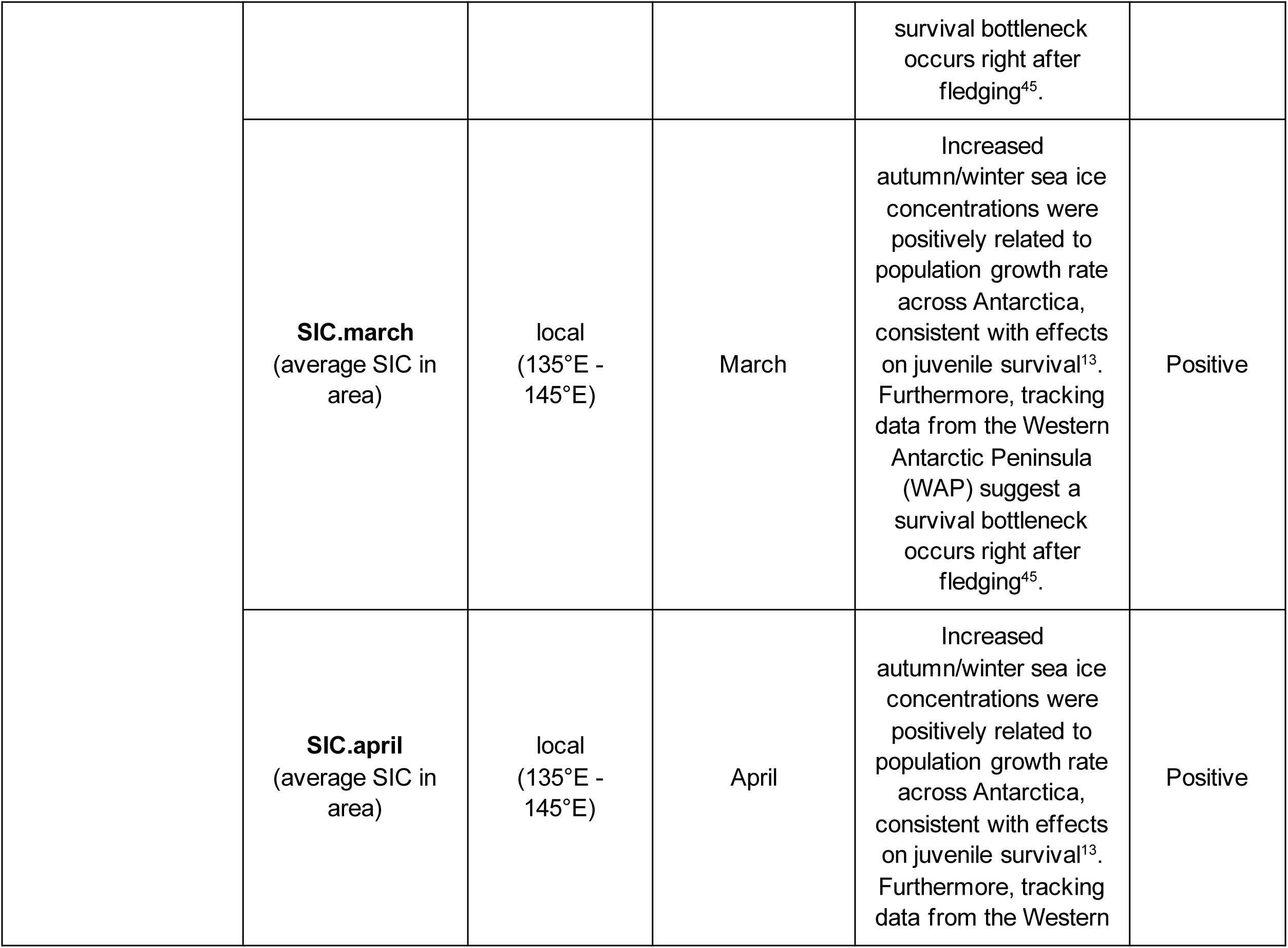
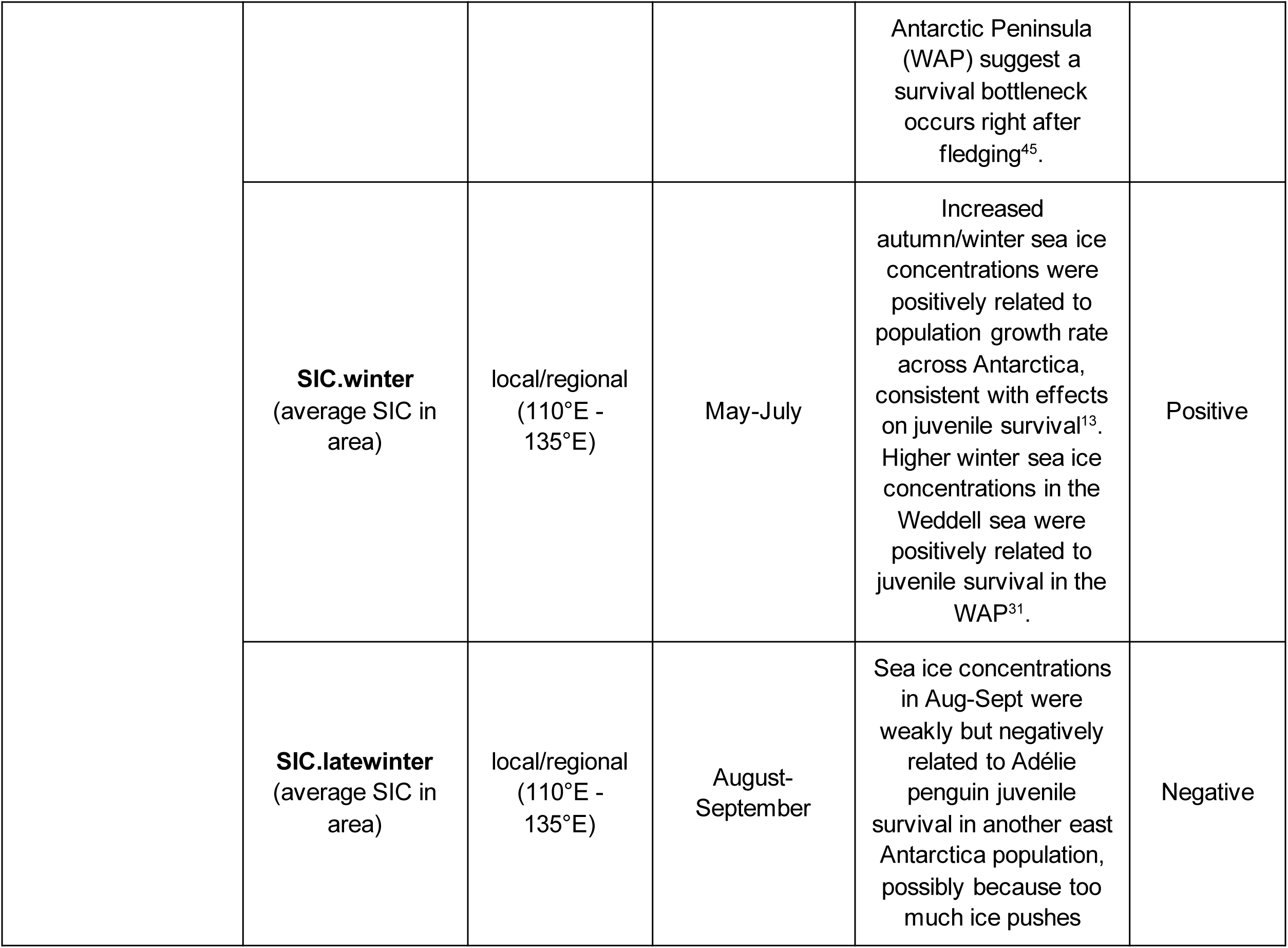
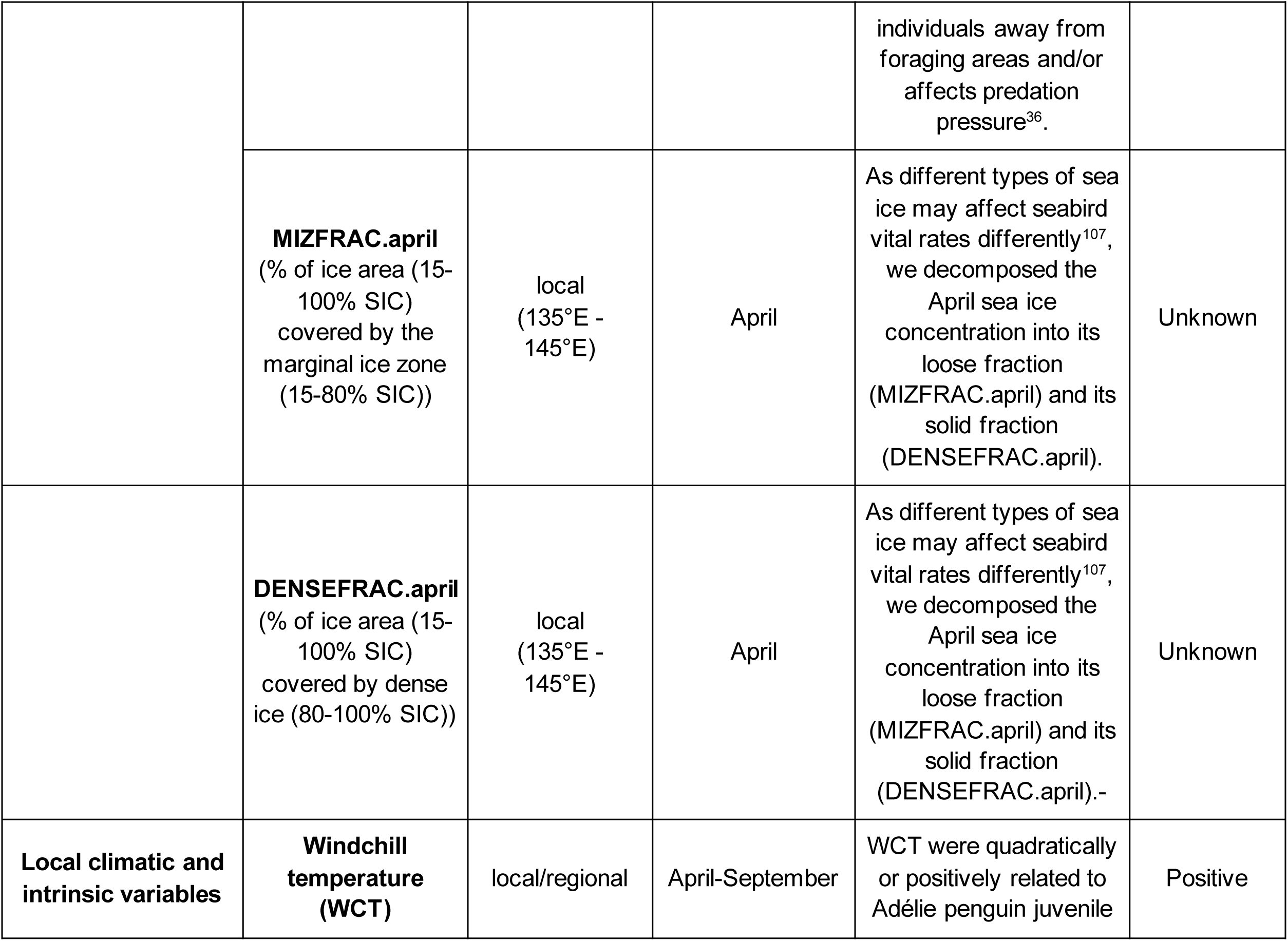
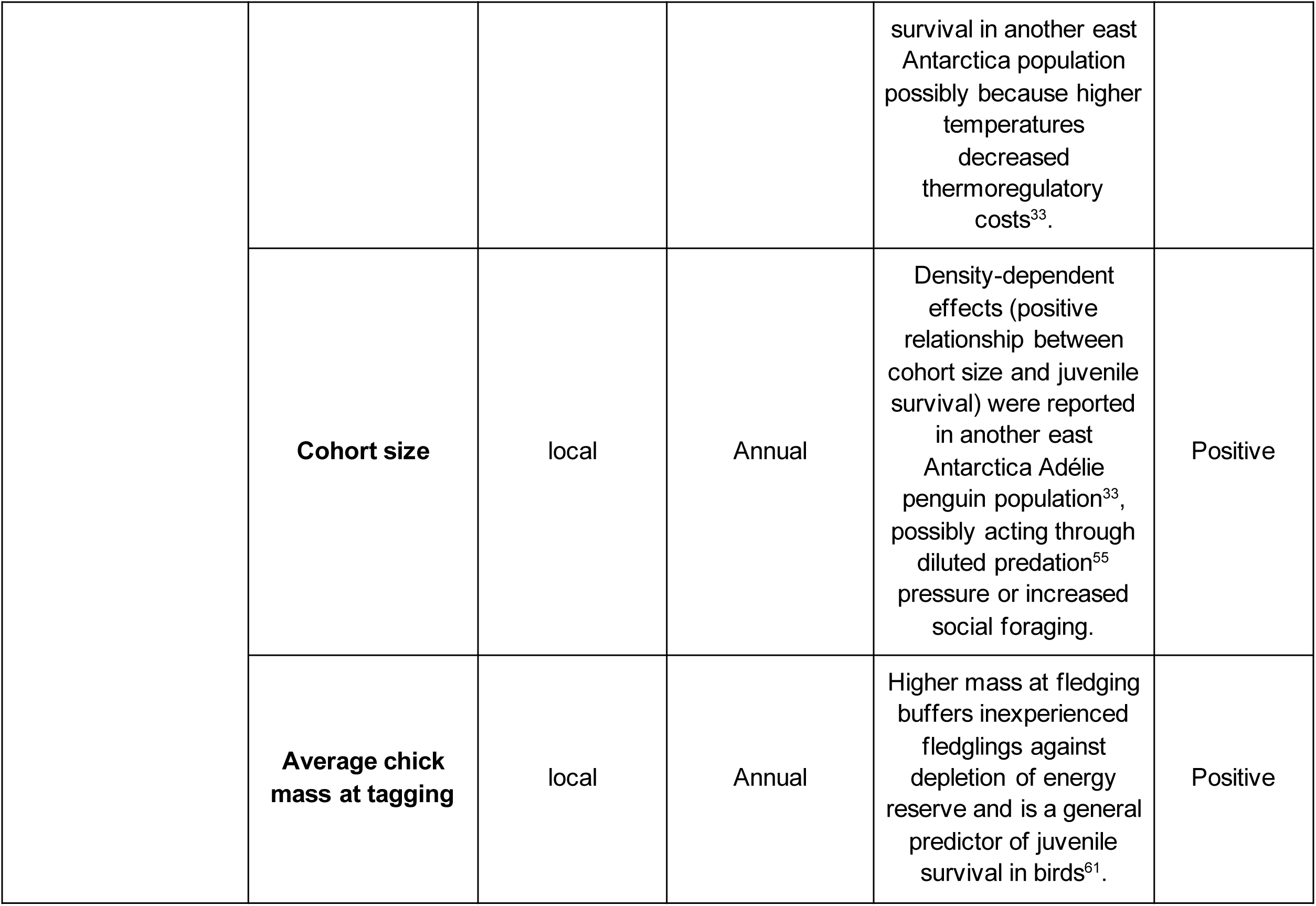
Annual covariates considered in the present study for explaining temporal variation in Adélie penguin juvenile survival.

**Table S3:** Model selection for recapture, transition, and survival probabilities. Abbreviations: AICc, Akaike Information Criterion corrected for small sample sizes; ΔAICc, AICc difference between current model and time-dependent model; K, number of parameters; Dev, Model deviance.

| Submodel set | Index | Model | AICc | $\Delta AICc$ (for each submodel set) | AICc Weights | K | Dev |
| --- | --- | --- | --- | --- | --- | --- | --- |
| *Recapture (p)* |  |  |  |  |  |  |  |
| | 3 | $\Phi$ (5 age classes × time) **p(state1: 3 age classes × time; state2: 3 age classes)** $\Psi$(5 age classes × | 12076.5 | 0.0 | 0.902 | 104 | 2276.1 |
| | 2 | $\Phi$ (5 age classes × time) **p(state1: 4 age classes × time; state2: 4 age classes)** $\Psi$(5 age classes × | 12081.8 | 5.3 | 0.063 | 114 | 2260.8 |
| | 1 | $\Phi$ (5 age classes × time) **p(state1: 5 age classes × time; state2: 5 age classes)** $\Psi$(5 age classes × | 12083.0 | 6.5 | 0.035 | 123 | 2243.4 |
| | 5 | $\Phi$ (5 age classes × time) **p(state1: 5 age classes + time; state2: 5 age classes)** $\Psi$(5 age classes × | 12095.7 | 19.2 | 0.000 | 74 | 2356.7 |
| | 6 | $\Phi$ (5 age classes × time) **p(state1: 4 age classes + time; state2: 4 age classes)** $\Psi$(5 age classes × | 12098.2 | 21.7 | 0.000 | 73 | 2361.3 |
| | 4 | $\Phi$ (5 age classes × time) **p(state1: 2 age classes + time; state2: 2 age classes)** $\Psi$(5 age classes × | 12135.6 | 59.1 | 0.000 | 93 | 2357.8 |
| *State transition ($\Psi$)* |  |  |  |  |  |  |  |
| | 3 | $\Phi$ (5 age classes × time) **p(state1: 3 age classes × time; state2: 3 age classes)** $\Psi$(5 age classes × | 12076.5 | 0.0 | 0.991 | 104 | 2276.1 |
| | 8 | $\Phi$ (5 age classes × time) **p(state1: 3 age classes × time; state2: 3 age classes)** $\Psi$(3 age classes × | 12086.9 | 10.3 | 0.006 | 100 | 2294.7 |
| | 7 | $\Phi$ (5 age classes × time) **p(state1: 3 age classes × time; state2: 3 age classes)** $\Psi$(4 age classes × | 12087.7 | 11.2 | 0.004 | 102 | 2291.4 |
| | 9 | $\Phi$ (5 age classes × time) **p(state1: 3 age classes × time; state2: 3 age classes)** $\Psi$(2 age classes × | 12096.4 | 19.9 | 0.000 | 98 | 2308.3 |
| | 10 | $\Phi$ (5 age classes × time) **p(state1: 3 age classes × time; state2: 3 age classes)** $\Psi$(state) | 12098.0 | 21.4 | 0.000 | 97 | 2311.9 |
| *Survival ($\Phi$)* |  |  |  |  |  |  |  |
| | 13 | **$\Phi$ (2 age classes × time)** **p(state1: 3 age classes × time; state2: 3 age classes)** $\Psi$(5 age classes × | 12044.7 | 0.0 | 0.997 | 75 | 2303.7 |
| | 12 | **$\Phi$ (3 age classes × time)** **p(state1: 3 age classes × time; state2: 3 age classes)** $\Psi$(5 age classes × | 12056.1 | 11.4 | 0.003 | 85 | 2294.7 |
| | 11 | **$\Phi$ (4 age classes × time)** **p(state1: 3 age classes × time; state2: 3 age classes)** $\Psi$(5 age classes × | 12064.6 | 19.9 | 0.000 | 95 | 2282.7 |
| | 3 | **$\Phi$ (5 age classes × time)** **p(state1: 3 age classes × time; state2: 3 age classes)** $\Psi$(5 age classes × | 12076.5 | 31.8 | 0.000 | 104 | 2276.1 |
| | 16 | **$\Phi$ (2 age classes + time)** **p(state1: 3 age classes × time; state2: 3 age classes)** $\Psi$(5 age classes × | 12127.1 | 82.3 | 0.000 | 65 | 2406.4 |
| | 15 | **$\Phi$ (2 age classes: cst/time)** **p(state1: 3 age classes × time; state2: 3 age classes)** $\Psi$(5 age classes × | 12264.6 | 219.8 | 0.000 | 68 | 2537.9 |
| | 14 | **$\Phi$ (5 age classes)** **p(state1: 3 age classes × time; state2: 3 age classes)** $\Psi$(5 age classes × state) | 12280.2 | 235.5 | 0.000 | 55 | 2580.0 |

**Table S4:**
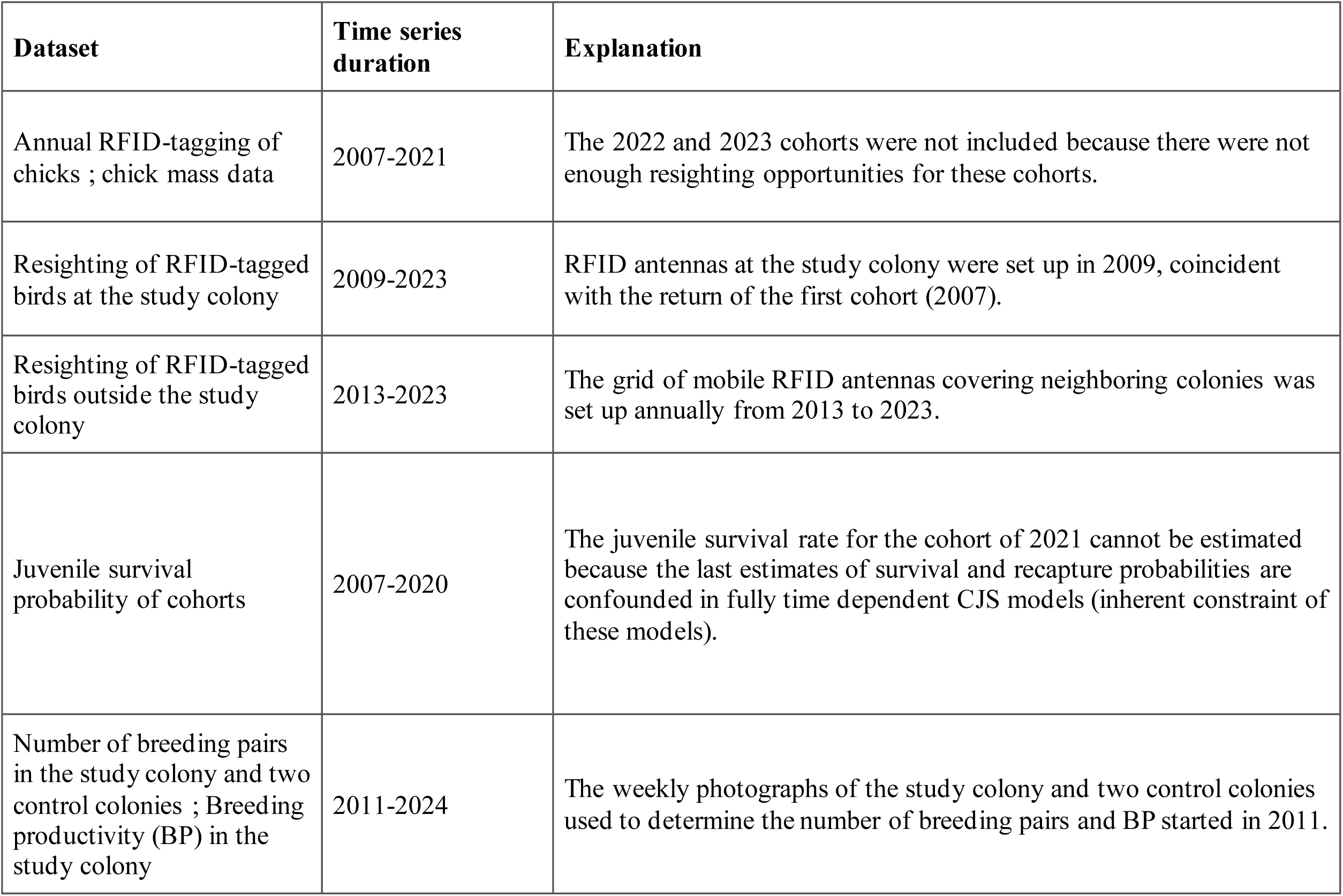
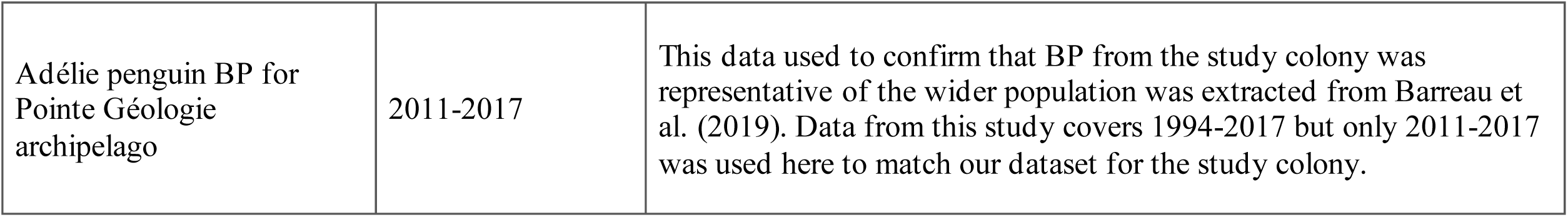
Timeline summary for the datasets used in the study. (datasets concerning the study colony only).

